# Identification of new insect Host Defense Peptides (HDP) from Dung Beetles (Coleoptera: Scarabaeidae) transcriptomes

**DOI:** 10.1101/2020.06.15.152363

**Authors:** Germán Alberto Téllez Ramirez, Juan Felipe Osorio-Méndez, Diana Carolina Henao Arias, Lily Johanna Toro S, Juliana Franco Castrillón, Maribel Rojas-Montoya, Jhon Carlos Castaño Osorio

## Abstract

**Introduction:** The Coleoptera Scarabaeidae family is one of the most diverse groups of insects on the planet, living in complex microbiological environments. Their immune systems have evolved with the generation of host defense peptides but only a small number of these peptides have been characterized.

**Methods:** In this work two sources of information were retrieved: 1) *De novo* transcriptomic data from two species of neotropical Scarabidae (*Dichotomius satanas* and *Ontophagus curvicornis*); 2) Sequence data deposited in available databases. A BLAST-based search against the transcriptomes with a subset of sequences representative of the Host Defense Peptides (HDP). The HDP were described with the cecropin, defensin, attacin, and coleoptericin families; their physical/chemical and structural properties were described.

**Results:** This work reports 155 novel sequences of HDP identified in 9 transcriptomes from seven species from the Coleoptera order: *D. satanas* (n= 76; 49.03%), *O. curvicornis* (n= 23; 14.83%), *T. dichotomus* (n= 18; 11.61%), *O. nigriventris* (n= 10; 6.45%), *Heterochelus sp* (n= 6; 3.87%), *O. conspicillatum* (n= 18; 11.61%) and *P. japonica* (n= 4; 2.58%). These sequences were identified based on similarity to known HDP insect families. New members of defensins (n= 58; 37.42%), cecropins (n= 18; 11.61%), attancins (n= 41; 26.45%) and coleoptericins (n= 38; 24.52%), with their physicochemical, structural characteristics, and sequence relationship to other insect HDP were analyzed.

**Conclusions:** 155 new HDP could be identified, based on similarity to known HDP insect families on nine transcriptome sequences of seven beetle species.

**Highlights:**

- This work identified 155 novel sequences of HDP found in nine transcriptomes from seven Coleoptera species.
- *De novo* transcriptomic data from two species of neotropical Scarabaeidae (*Dichotomius satanas* and *Ontophagus curvicornis*).
- *In silico* prediction of physicochemical properties, structural features, sequence similarity, and antimicrobial activity of Scarabaeidae HDP.

## Introduction

One of the main effectors of the insect immune response is the production of Host Defense Peptides (HDP) or antimicrobial peptides. Families of these peptides have been identified in all taxonomic groups, thus representing an ancient and efficient defense mechanism against pathogens. In insects, most HDP are synthesized as precursors or pro-proteins in the fat body and hemocytes [1–3]. Host Defense Peptides are cationic, amphipathic polypeptides, produced in all known genera of living organisms, and represent an ancient innate defense mechanism [4–6]. Once activated by post-translational proteolysis [1,7–9], they act as effector molecules against pathogens in a broad spectrum of antimicrobial activity against Gram-positive and Gram-negative bacteria, protozoa, fungi, and viruses. They also have low propensity for developing resistance. This efficiency is thought to be one of the biological attributes that would explain the evolutionary success of insects [10,11]. Therefore, they have received attention for the development of new antimicrobials with clinical applications in recent years [12,13].

Insect HDP have been classified according to their sequence, physicochemical and structural properties, in cecropin, defensin, attacin, and coleoptericin families. Other families, such as moricin and gloverin have been identified only in Lepidoptera [2,14,15]. Cecropins are a family of 3–4 kDa, cationic, alpha helix, amphipathic peptides devoid of cysteine residues [16,17]. Mature active cecropins are generated after removal of the signal peptide and form two α-helices connected by a hinge. A long hydrophobic C-terminal and a strongly basic N-terminal domain is presumptively required for the biological activity and they are most active against Gram-negative bacteria [18–22].

Defensins are the largest family of HDP and are ubiquitous in almost all forms of life, including animals, fungi, and plants [23]. They have broad-spectrum antimicrobial activity against bacteria, fungi, and viruses [24]. The majority of mature defensins are cationic peptides composed of 24 to 42 amino acid residues, characterized by six cysteines [25]. The structure of insect defensins is composed of an α-helix, followed by an antiparallel β-sheet linked by three intramolecular disulfide bonds, forming a “Cysteine-Stabilized alpha Beta (CSαβ)” or “loop-helix-beta-sheet” structure [26,27].

Attacins are larger peptides with basic (~ 8.3) or acidic (~ 5.7) isoelectric points, and molecular weights of 20-23 kDa; we can find two isoforms, basic and acids attacins [28]. Their secondary structure is composed of a hydrophobic alpha helix similar to glycine-rich peptides [29]. They can inhibit growth of Gram-negative bacteria, and the synthesis of bacterial proteins like OmpC, OmpF, OmpA and LamB [30,31].

Coleoptericins contain approximately 70 amino acid residues and are characterized as glycine and proline-rich antimicrobial peptides that are bactericidal against Gram-positive and Gram-negative bacteria [32]. There are two subgroups, one positively and other negatively charged, and their C-terminus has a basic nature [32,33].

With approximately one-million characterized species, insects represent the largest class within the animal kingdom and Coleoptera are the most diverse order and is the most diverse in insects [34–36]. Nevertheless, only 305 of the 3070 HDP sequences deposited in the Antimicrobial Peptide Database (APD) are derived from insects [37]. To date some HDP have been described in the Coleoptera order, as is the case of *Allomyrina dichotoma* [38], *Octodonta nipae* [39], *Hylobius abietis* [40], *Nicrophorus vespilloides* [41], *Tenebrio molitor* [42], *Calomera littoralis* [43], *Protaetia brevitarsis seulensis* [44], *Tribolium castaneum* [45–47], *Holotrichia diomphalia* [48], *Zophobas atratus* [33], *Allomyrina dichotoma* [32], *Acalolepta luxuriosa* [49], and *Sitophilus oryzae* [50]. The HDP reported in the Scarabaeidae family are scarce compared with their wide diversity of species consisting of over 30.000 globally. The sequence and function of only a few of these peptides have been characterized, including those of the Coprisin, Oxysterlin, and Scarabaeicin families [51–57]. Therefore, we propose to identify and describe new putative HDP in publicly available assembled transcript sequences from the NCBI Transcriptome Shotgun Assembly (TSA) database of seven different species of Scarabaeidae and two new transcriptomes from the neotropical beetles *Dichotomius satanas* and *Ontophagus curvicornis*, both species widely distributed inhabiting the Andean region of Colombia [58–61].

## Materials and Methods

### Collection and maintenance of beetles

Neotropical dung beetles used in this research were obtained in the municipality of Filandia, Quindío-Colombia (4.686998” N and −75.614500” W; datum=WGS84) 1.923 m.a.s.l. The beetles captured were identified as *Dichotomius satanas* and *Ontophagus curvicornis* with the Cultid-Medina, 2012 taxonomic key [58]. Once collected, they were maintained in a terrarium with organic soil and human feces bait for 12 h. Then, they were separated into two groups of five individuals each. One group was inoculated in the ventral lateral abdomen with 10 μL of a pool of 1X10^6^ UFC/mL formalin-fixed bacteria (*E.coli* and *Staphylococcus aureus*) and fungi (*Candida albicans*). The other group was used as an untreated control. Finally, the fat body and part of the hindgut were dissected 12 h post-inoculation.

### Total RNA extraction, transcriptome sequencing, and *de novo* assembly

Total RNA was extracted by using Ambion total RNA extraction kit (Invitrogen cat PureLink^®^ RNA Mini Kit, life technologies 12183025). Total RNA was prepared by using bead clean up and library preparation with Illumina RNA poly-A selection. The transcriptome was sequenced with Illumina Hiseq and pair-end read with 150bd length. The FASTAq files were checked by FastQC [62], the trimming was done by Trimmomatic V0.36 [63], the data from each species were merged and the transcriptome was *de novo* assembled by using Trinity V2.5. on Indiana University National center for genome analysis support (https://galaxy.ncgas-trinity.indiana.edu/) [64].

### Transcriptome shotgun assemblies

Transcriptome shotgun assemblies (TSA) from Scarabaeidae species: *Trypoxylus dichotomus, Onthophagus nigriventris, Onthophagus curvicornis, Popillia japonica, Heterochelus sp, Dichotomius satanas*, and *Oxysternon conspicillatum* were downloaded from the sequence set browser (https://www.ncbi.nlm.nih.gov/Traces/wgs/). The fasta files of the assembled transcriptomes were converted into BLAST databases with CLC main workbench software 7.9.1.

### Homology identification of HDP

The queries from the Cecropin family were constructed from Oxysterlins [53] and different sequences related to the HDP InterPro families (Cecropin: IPR020400; Defensin: IPR017982; Coleoptericin: IPR009382, and Attacin: IPR005520 IPR005521) and a multi-TBLASTn search was constructed and the result sequences were filtered according to the E-score ≤ 0.01 and the open reading frame (ORF) was identified for each sequence [65]. The list of sequence codes identified in the tBLASTn was extracted from the different transcriptomes and the related amino-acid sequence was identified by using the ORF finder tool (https://www.ncbi.nlm.nih.gov/orffinder/) [66].

### Workflow to analyze putative HDP sequences

The presence and location of signal peptides in the deduced HDP amino acid sequences were predicted with SignalP (**http://www.cbs.dtu.dk/services/SignalP/**) [67]. The physicochemical characteristics of the peptides (molecular mass, isoelectric point, and Kyte-Doolittle hydrophobic profile) were calculated with the protein report tool from the CLC main workbench V7.9.1. The total net charge was calculated with the APD3 (**http://aps.unmc.edu/AP/main.php**) calculation and prediction tool [37]. Prediction of the antimicrobial function was conducted with the SVMC, RFC, and DAC algorithms available in Campr3 [68], Classamp [69], and iAmppred tools [37].

### Structural analysis

The secondary structure prediction was done with the Psipred (**http://bioinf.cs.ucl.ac.uk/psipred/**) server [70]. Prediction of functional domains was done with Interpro (https://www.ebi.ac.uk/interpro/) [71]. The tertiary structure was predicted with RAPTOR X (http://raptorx.uchicago.edu/StructurePrediction/predict/) [72], and structural alignments were done with the 3Dcomb V1.18 tool [73]. All the models were visualized in UCSF Chimera V1.13.1. [74]. The protein-protein interactions of the sequences modeled were constructed by using the Cluspro server [75]. Illustration of the general structural properties for each family was made by DOG V1.0 [76].

### Similarity dendrogram

The sequences of InterPro families (cecropin: IPR020400; insect Defensin: IPR017982; Coleoptericin: IPR009382 Attacin: IPR005520 IPR005521) and the taxonomic key corresponding to each sequence were downloaded from the PIR batch server (https://pir.georgetown.edu/pirwww/search/batch.shtml). The signal and propeptide were identified and removed and the mature peptides were aligned with the HDP from the Scarabaeidae with Clustal Omega (https://www.ebi.ac.uk/Tools/msa/clustalo/) or MUSCLE [77]. Dendrograms were generated by Neighbor-joining with Jukes-Cantor protein distance and bootstrap with 10000 replicates in CLC main workbench V7.9.1.

### Ethics statement

This work was approved by the Universidad del Quindío bioethics committee under act number 8 of 06 May 2016. The contract of access to genetic resources was drawn through resolution N° 120 of 22 October 2015 with the Colombian Ministry of Environment and Sustainable Development.

## Results and Discussion

### Transcriptome shotgun assemblies (TSA)

Traditional strategies for HPD identification and characterization involve biochemical purification methods with RP-HPLC (reverse-phase high-pressure liquid chromatography) coupled with mass spectrometry and functional assays. Other strategies use highly conserved positions of some HDP families to identify potential HDP by similarity searches or molecular biology approaches with RACE-PCR (rapid amplification of cDNA ends) [78]. Artificial neural network algorithms have also been trained with information about structural and physicochemical characteristics to identify novel HDP sequences with proteomic and genomic methodologies [79]. Recent developments in high-throughput sequencing technologies have represented a novel and efficient method for gene identification [80]. Transcriptome-based approaches using next-generation sequencing are particularly useful because they focus on the expressed (*i.e*. exomic) portion of the genome. This information can be exploited by using bioinformatic tools to search for target genes/proteins using a step-by-step selection strategy [81]. In this work, the transcriptomes of two Scarabaeidae species (*O. curvicornis* and *D. satanas*) were sequenced, assembled and submitted to the DDB/EMBL/GenBank database under the accession codes GHMD00000000 - GHMA00000000, bio-sample: SAMN10614998 - SAMN10614917 and bio-project: PRJNA510790 - PRJNA510790, respectively. The transcriptome characteristics used for this work are shown in **Table 1**.

**Table 1.** TSA from Scarabaeidae.

| Prefix (TSA code) | Organism | Bio project code | Biosample code | Source | Contigs number |
| --- | --- | --- | --- | --- | --- |
| GAQV01 | *Trypoxylus dichotomus* | PRJNA231720 | SAMN02444008<br>SAMN02444009<br>SAMN02444010<br>SAMN02444011<br>SAMN02444012<br>SAMN02444013<br>SAMN02444014<br>SAMN02444015<br>SAMN02444016<br>SAMN02444017<br>SAMN02444018<br>SAMN02444019 | 3rd instar larval<br>and prepupal | 34455 |
| IABQ01 | *Trypoxylus dichotomus tunobosonis* | PRJDB4830 | SAMD00051587 | 3rd larvae | 30157 |
| GAQW01 | *Onthophagus nigriventris* | PRJNA231725 | SAMN02444020<br>SAMN02444021<br>SAMN02444022<br>SAMN02444023 | 3rd instar larval<br>and prepupal | 59302 |
| GARJ01 | *Popillia japonica* | PRJNA233626 | SAMN02569976 | Antenna | 698 |
| GARK01 | *Popillia japonica* | PRJNA198730 | SAMN02055564 | Grub | 1916 |
| GDNJ01 | *Heterochelus*<br>*sp. AD-2015* | PRJNA286531 | SAMN03799575 | Tissue | 50435 |
| GEXM01 | *Oxysternon conspicillatum* | PRJNA339294 | SAMN05589108 | Adult fat body | 27567 |
| GHMA01 | *Dichotomius satanas* | PRJNA510790 | SAMN10614917 | Adult fat body<br>and intestine | 463430 |
| GHMD01 | *Onthophagus curvicornis* | PRJNA510790 | SAMN10614998 | Adult fat body and intestine | 172518 |

A tBLAStn search on the nine assembled transcriptomes was done with the constructed HDP sequence queries, the result contigs were retrieved and only those containing complete ORF were considered for further analysis. In total, 155 contigs encoding for potential HDP were identified. The number of HDP identified correlated with the size of the transcriptome and the condition of the sample (**Fig 1A**). These three species, *O. curvicornis*, *D. satanas*, and *O. conspicillatum*, have most of the HDP. This may be explained by the fact that these three specimens had been obtained from the fat body of adult specimens previously subjected to stimulation to boost the insect immune response; these results agree with the different expression patterns of HDP with basal and inducible forms in different tissues and conditions [2,82].

**Fig 1.**
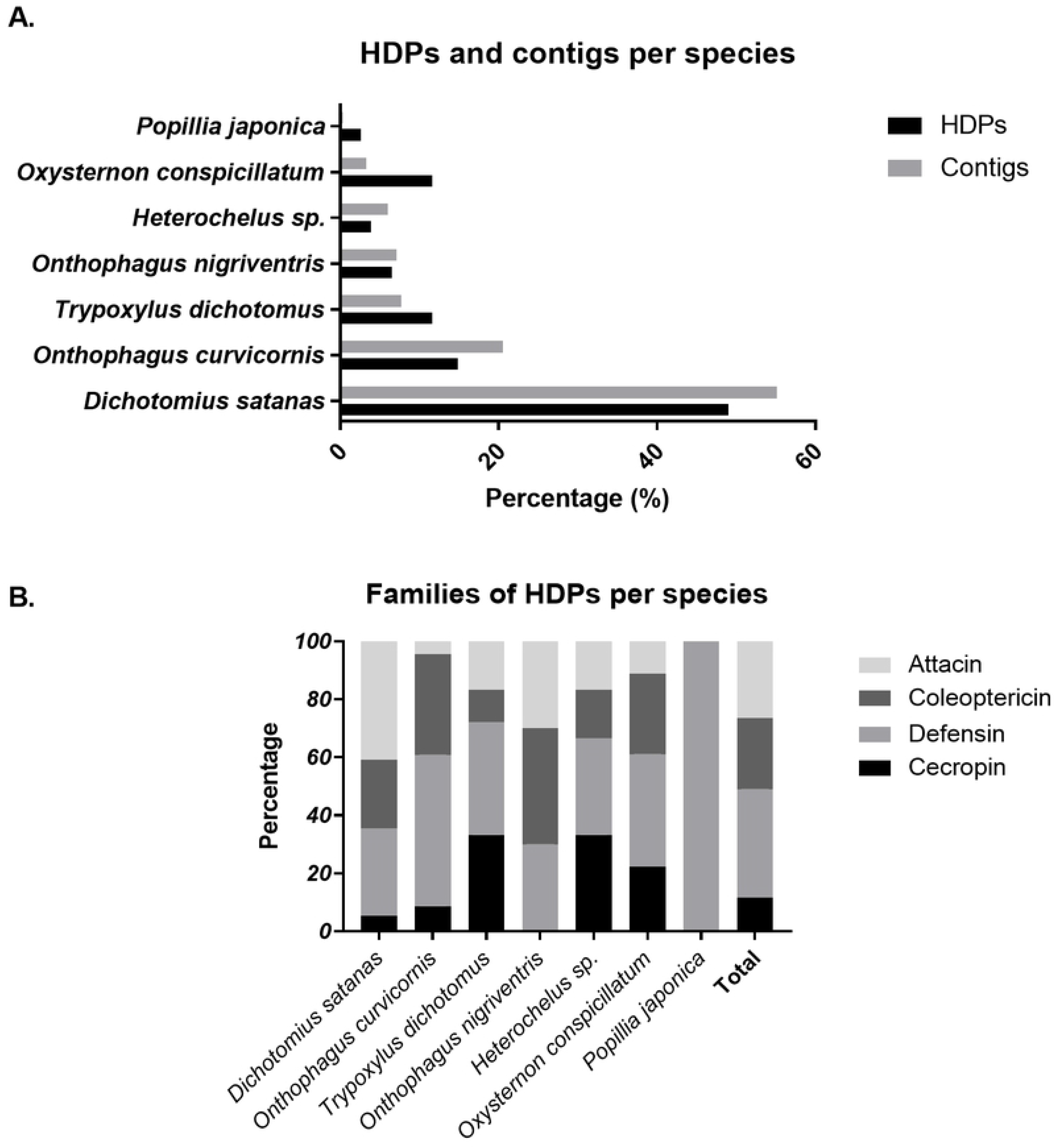
Percentage of contigs in nine Scarabaeidae TSA and percentage of HDP identified by sequence homology. **A**. Percentage of contigs and HDP per species. **B**. Percentage of Scarabaeidae HDP families by dung beetle species.

Four HDP family distribution also varied according to species (**Fig 1B**) with members from these families identified in all species, with two exceptions *Popillia japonica* with only defensin peptides and *Onthophagus nigriventris* where the cecropin family was missing. *Heterochelus* sp and *O. nigriventris* have less number of defensins in contrast with *O. conspicillatum, O. curvicornis, T. dichotomus*, and *D. satanas* wich have more attacins. The defensin HDP family has the majority of peptides representing 37.42 % (**Fig 1B**). The difference in the number of HDP found among species and type of HDP may be explained by the different tissues sampled, size of the transcriptome, and life cycle stage of the individual sample, according to these differences, it has been described that the expression of HDP is influenced by context-specific characteristics, where sex, presence of offspring, and carcass affect their expression in a complex system of transcriptional reprogramming, reflecting adaptations to specific ecological niches [82].

This article, Identified 155 new putative HDP sequences from nine transcriptomes (**S1 appendix**), while another work with a similar approach using tBLASTn and BLASTp searches in newly sequenced arthropod genomes and expressed sequence tags (EST) derived from the red flour beetle (*Tribolium castaneum*), monarch butterfly (*Danaus plexippus*) and human body louse (*Pediculus humanus humanus*) identified six HDP [83]. One difference in the results may be explained by the use of the TSA database where the sequences may be more refined in the context of genome complexity with different introns and gene arrangements; another difference was the query construction, where sequences reported in APD were used and our work used a greater query search according to the InterPro related sequences.

### Cecropin family description

Cecropins are the most abundant family of linear α-helical HDP in insects [84]. They have been identified in Hexapoda orders like Coleoptera, Diptera, and Lepidoptera [26]. Insect and non-insect cecropins are encoded by non-homologous genes. However, in holometabolous orders they have evolved only once [85]. Genes from this family frequently encode for tryptophans at the first or second position of the mature peptide and lack cysteines. They are basic amphipathic peptides rich in cationic amino acids, with an N-terminal hydrophilic domain and a C-terminal hydrophobic domain. Structurally, they are characterized by two distinctive alpha-helical segments linked by a short hinge. In the post-translational level, they have a signal peptide and a frequent C-terminal amidation [16,17,19,26,86].

In Scarabaeidae, 18 cecropin sequences were found with mature peptide lengths ranging from 37 to 55 residues, with molecular weights around 4 kDa. The sequence alignment shows a highly conserved signal peptide, the mature peptide has an N-terminal cationic domain [GR]-[SW]-K-[RKG]-[WLF]-R-K-[FIL]-E-[KR]-[RKA]-[VSG]-[KR]-[KR] with high frequency of K-R residues and hydrophobic angle from 120° to 180°. The C-terminal domain has a higher degree of variability with a region rich in acid residues and an aliphatic hydrophobic region (**Fig 2**). The predicted secondary and tertiary structures show highly conserved alpha-helix with a TM score of 0.51 in the structural alignment, suggesting that they can be classified as a single structural family (**Fig 3**) [73,87,88]. We found that only GEXM01014653.1 and GEXM01014651.1 sequences have an additional segment in the N-terminal domain, probably because of alternative splicing or a transcriptome assembly error.

**Fig 2.**
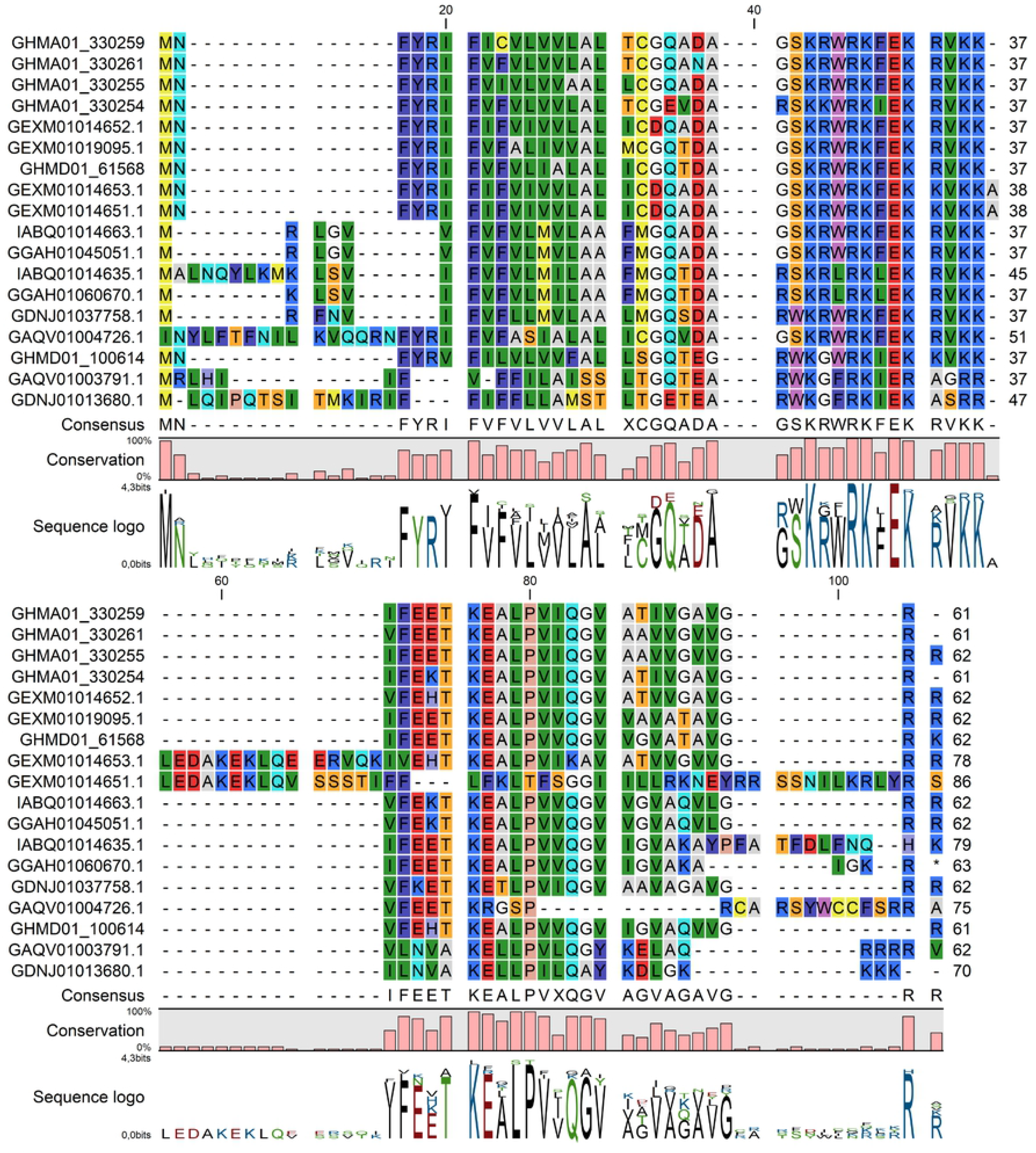
Multiple-sequence alignment of cecropin HDP found in Scarabaeidae. Position 40 in the alignment represents the cleavage site of the signal peptide predicted with SignalP.

**Fig 3.**
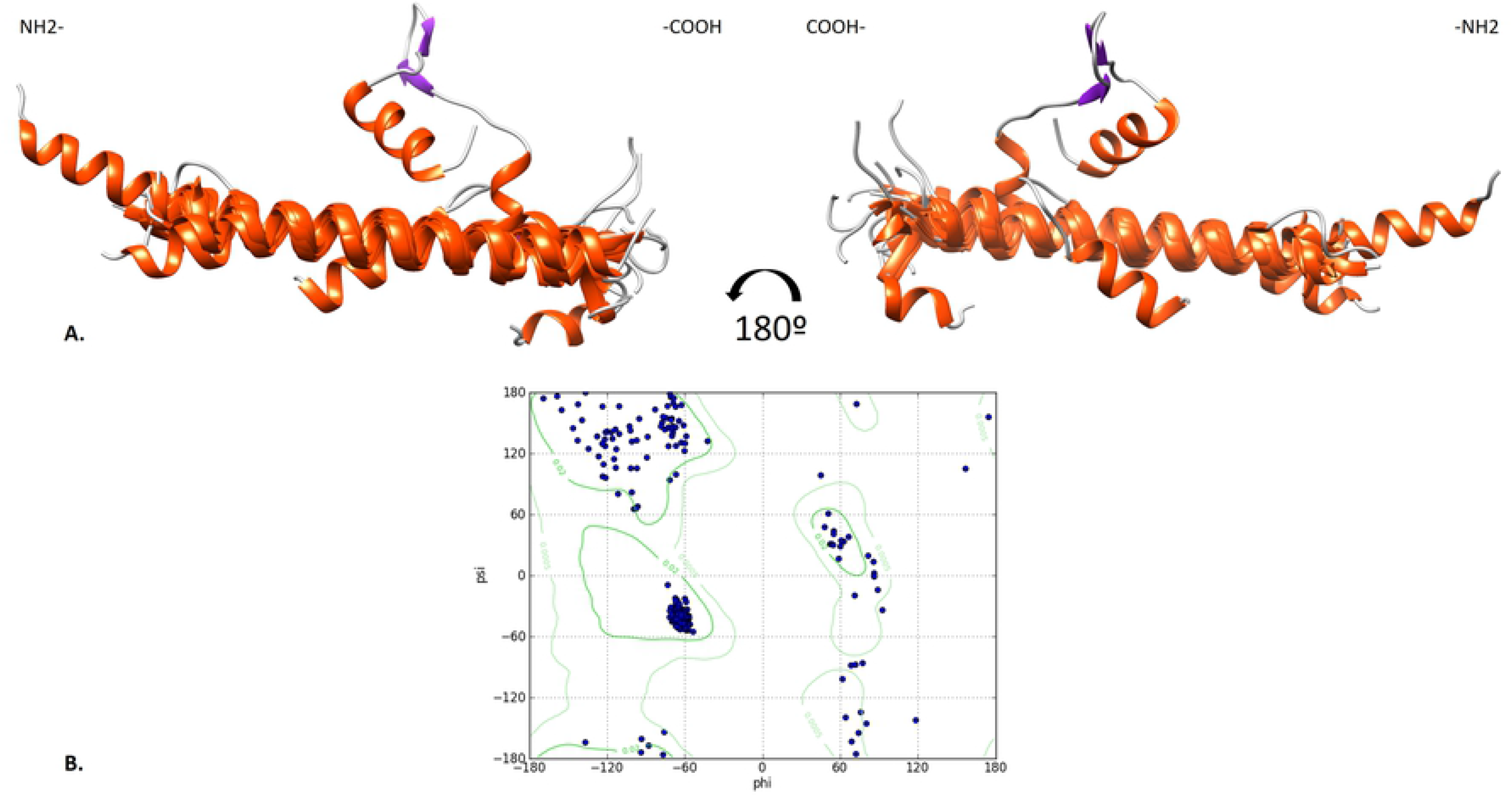
Structure of Scarabaeidae mature cecropins. **A**. Structural superposition constructed in DeepAlign. TMscore: 0.517 with Alpha helical conserved tertiary conformation. **B**. Ramachandran plot of the superimposed structures.

The insect cecropin dendrogram shows that the majority of sequences are from Diptera and Lepidoptera orders as already described [85]. The dendrogram structure has four main clades that are well related to the phylogenetic orders, a highly conserved Diptera clade representing flies, Lepidoptera, Ascaridida, and a distant share clade of Coleoptera and mosquitoes (**Fig 4**). Compared to other orders like Diptera and Lepidoptera, cecropins from Scarabaeidae show more variation and divergence among the sequences within the same order.

**Fig 4.**
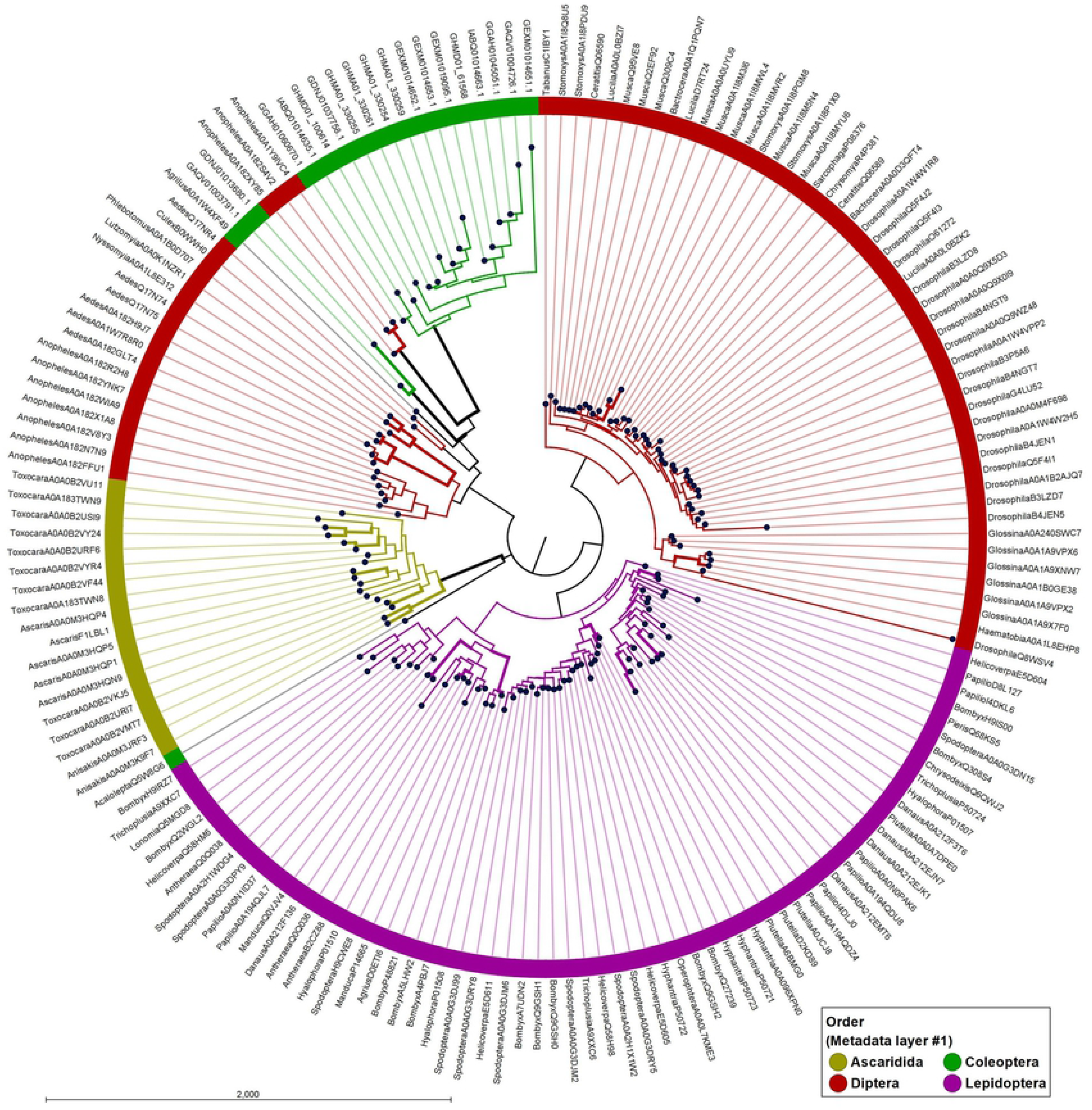
Neighbor-joining similarity dendrogram of invertebrate cecropins. Sequences from the mature peptides were aligned by MUSCLE. The taxonomic distribution of the sequences is indicated in the presented color code. Confidence values of the branches were calculated with 10.000 bootstrap replicates. Thicker lines show branches with bootstrap threshold value > 70.

According to the previously described cecropins in Coleoptera, there are few representatives with Oxysterlins (1, 2, and 3) (*O. conspicillatum*), Cec (*Acalolepta luxuriosa*), and Sarcotoxin Pd (*Paederus dermatitis*) [53,89,90]. Interestingly, the Interpro data set, also has low representation of Coleopteran cecropins, with only two representative sequences, *Acalolepta luxuriosa* (Q5W8G6) and *Agrilus planipennis* (A0A1W4XF49). Additionally, the InterPro search fails to classify the Scarabaeidae cecropin sequences as insect cecropins, indicating that, these sequences are different from the InterPro signatures; nevertheless, as described, these new peptides had the same physicochemical and structural characteristics of the insect cecropins [84].

### Defensin family description

Insect defensins are inducible antibacterial peptides with a spectrum of activity focused mainly against Gram-positive bacteria, although activity against fungi and parasites has also been reported [91,92]. They are extensively distributed in nearly all life forms [24]. In insects, they have been found in Diptera, Hymenoptera, Coleoptera, Trichoptera, Hemiptera, and Odonata orders [14,91]. Defensins are 18 - 45 amino acids long with 6 - 8 conserved cysteine residues stabilized by three or four disulfide bonds [92,93]. Typically, insect defensins have the same cysteine pairing: Cys1-Cys4, Cys2-Cys5, and Cys3-Cys6. Two disulfide bonds connect the C-terminal β-sheet and the α-helix; and the third connects the N-terminal loop with the second β-sheet. This structural topology is known as cysteine-stabilized αβ motif (CSαβ) and is common among defensin peptides across different organisms [34,94–96]. Once synthesized, pre-pro-defensins are proteolytically processed. First is the removal of the signal peptide to produce an inactive prodefensin; then, it suffers an additional cut of the propeptide by a furin-like enzyme in an R-X-[RK]-R conserved site to produce an active mature peptide [14,97].

This work identified 58 new defensin sequences in Scarabaeidae. As described for other insect defensins, those encoded by Scarabaeidae encode for a signal peptide followed by a propeptide (position 51-80 **Fig 5**) characterized by acid residues and an R-X-[RK]-R furin-like cleavage site (**Fig 5**). These sequences were classified into three groups (Group A, B, and C), according to their sequence, structural, and physicochemical properties.

**Fig 5.**
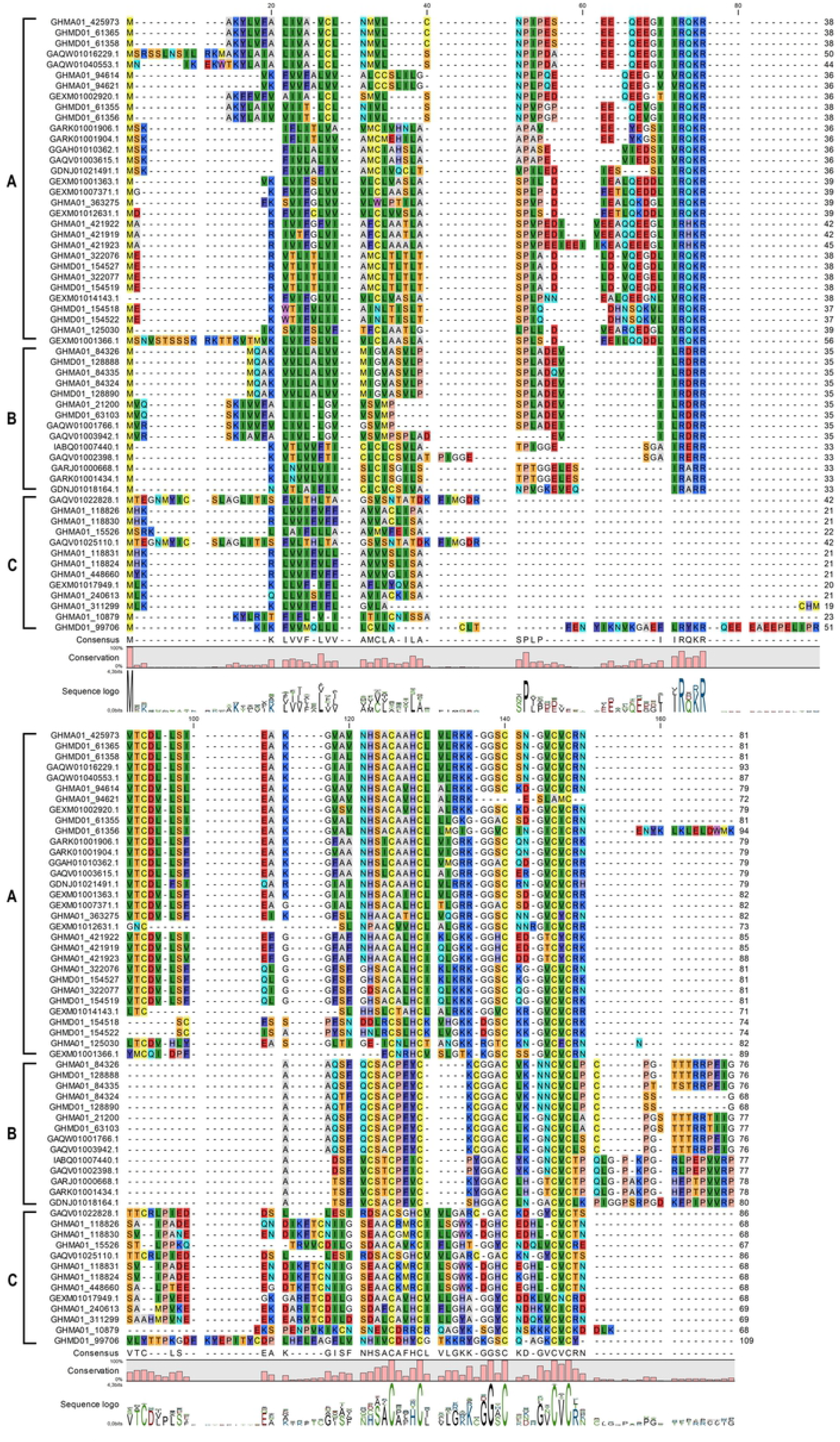
Multiple-sequence alignment of defensin HDP found in Scarabaeidae. The Scarabaeidae defensin groups are annotated in the left keys. The predicted signal peptide cleavage site is in position 50. Position 72-75 indicates the propeptide furin-like cleavage site.

Defensin group A with 31 sequences show a classical defensin pattern of helix beta-sheet structure with three disulfide bridges between cysteine pairs Cys1-Cys4, Cys2-Cys5, and Cys3-Cys6 (20 sequences). For 10 sequences the first cysteine pairing seems to be lost, but they keep the Cys2-Cys5 and Cys3-Cys6 binding pattern. There is only one sequence (GHMA01_94621) with no predicted disulfide bridges. Structurally, there is one subgroup within Group A that can be distinguished by the absence of the N-terminal hydrophobic loop (six sequences) (**Fig 6**).

**Fig 6.**
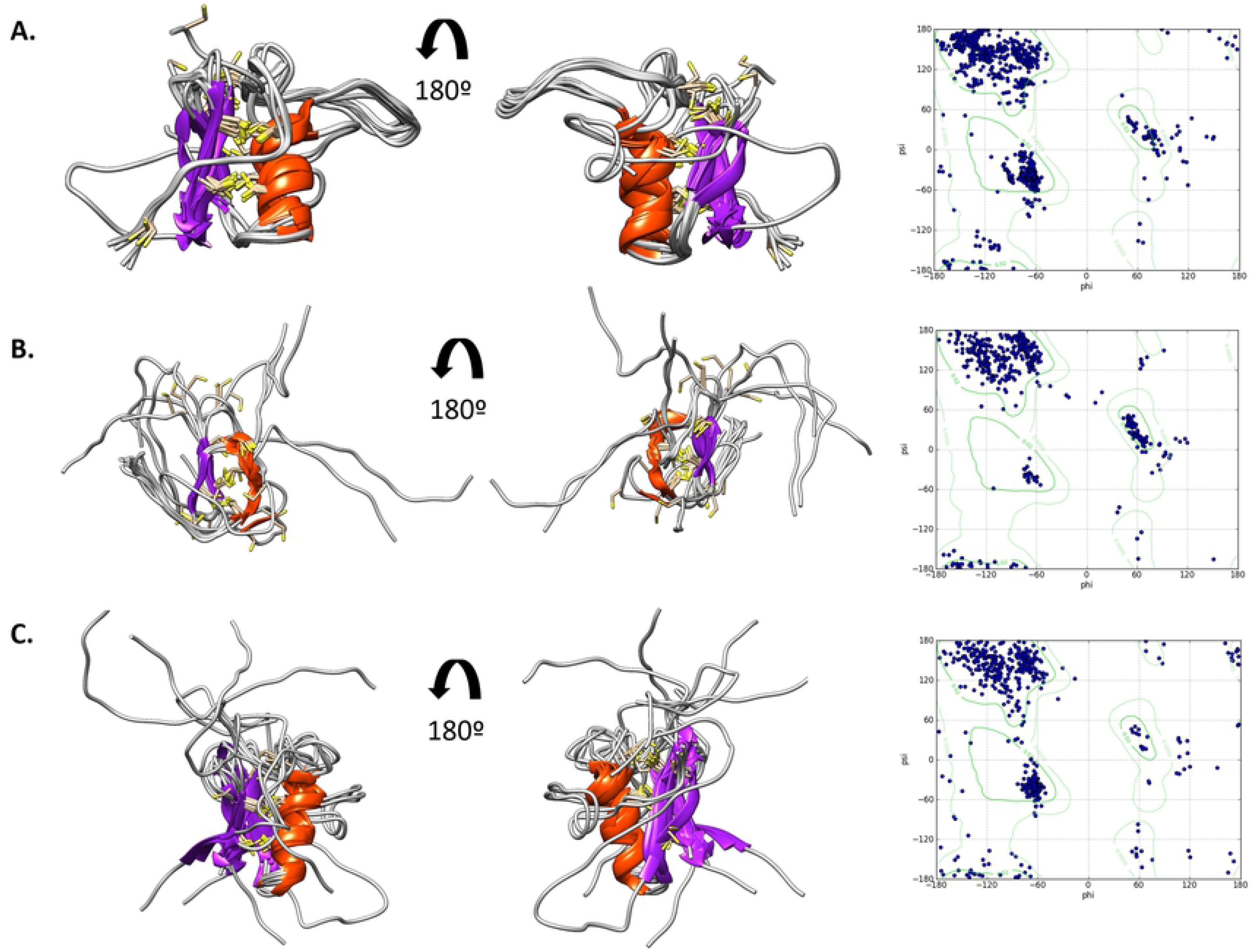
Structural superposition of the mature Scarabaeidae Defensins and Ramachandran plot, constructed in DeepAlign. **A** Group A with a helix beta-sheet structure (TM-score: 0.569). **B**. Group B with left-handed helix. and high frequency of residues in the left-handed helix region lying between 30° and 130° in the Φ angle and −50° and 100° in the ψ angle in the Ramachandran plot (TM-score: 0.571). **C**. Group C with helix beta-sheet structure (TM-score: 0.425).

Defensins group B is represented by 14 sequences. One key feature in this subgroup is the left-handed helix 6 to 8 residues long, related to a P-F-[YVI] motif in position 11. They encode for 6 or 8 cysteine residues (nine sequences) that form two predicted disulfide bridges between pairs Cys2-Cys4 and Cys3-Cys5 (six sequences); or Cys2-Cys5 and Cys3-Cys6 (six sequences). Two sequences (GHMA01_21200; GHMD01_63103) have one pair Cys1-Cys6 and loss predicted helix of the tertiary structure prediction. A hydrophobic random coil region was found in the C-terminal end of the sequences (**Fig 6**). The left-handed helix is a rare structural motif found in peptides and proteins. It has been found in regions related to protein stability, ligand binding, or as part of an enzyme’s active site. Thus, a significant structural or functional role for this secondary structure element has been suggested. The motif related to this particular structure agrees with the described propensity of amino acids to form such structure, preferring aromatic and large aliphatic amino acids [99,100]. To our knowledge, this kind of structure has not been described in the defensin family but its appearance may indicate functional importance due to its unique structural parameters.

Defensins group C with 13 sequences contains the classic three disulfide bridges of the insect defensins between cysteine pairs Cys1-Cys4, Cys2-Cys5, and Cys3-Cys6. Interestingly, they lost the R-X-[RK]-R cleavage site conserved for the other defensins, thus adding 15 N-terminal negatively charged residues to the mature peptide. In addition, there are two highly conserved acidic residues [DE] in 45 and 64 positions located at the beginning of the alpha-helix and beta-sheet loop (**Fig 6**). These acid residues explain the negative charge of this group (**Fig 7**). These types of anionic antimicrobial peptides have been shown to kill the human B-defensin-resistant Gram-positive bacterium *Staphylococcus aureus*, which escapes attacks from cationic peptides probably by incorporating positive charges on the membrane surface by adding Lys to lipids [98]. The anionic antimicrobial peptides, although rarely documented appear to complement the cationic antimicrobial peptides, offering a complete spectrum of antimicrobial peptides [99,100].

**Fig 7.**
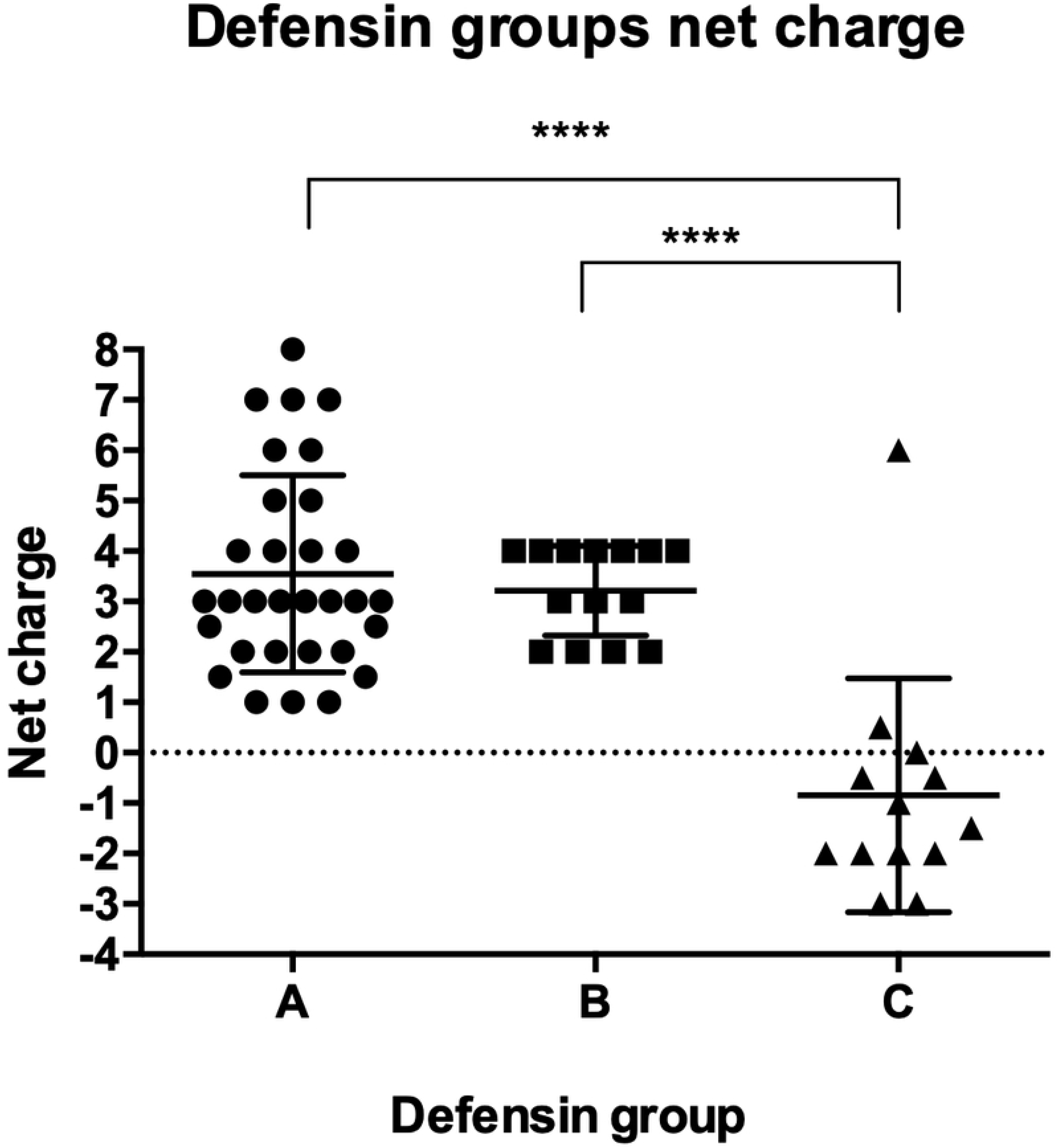
The net charge of the Scarabaeidae defensins predicted by APD. Statistical differences were evaluated by ANOVA test with Tukey’s multiple comparisons (p-value: ****<0.0001).

To evaluate the relationships of the Scarabaeidae defensins identified, a dendrogram was constructed with the retrieved insect defensins reported under the Interpro IPR017982. The sequences analyzed are distributed in the six major orders of insects (**Fig 8**). The distribution shows four main clades, two related to the Hymenoptera order, and one distinctive clade for Diptera and Hemiptera. Orders, like Coleoptera, Phthiraptera, and Archaeogastropoda were not grouped in a single clade, representing a more diverse distribution throughout the diversity of sequences. The defensins group A from Scarabaeidae was related to the Coleopteran defensin described from the Interpro. Group B of Scarabaeidae defensins were exclusively found in Scarabaeidae in a closer relationship with the clade corresponding to Hymenoptera. Group C seems to be related to a single defensin encoded by Hymenoptera (VespidaeXP_014602557) and other Coleoptera species. These results are compatible with the idea that groups B and C are Scarabaeidae-specific groups of defensins with novel structural and physicochemical properties.

**Fig 8.**
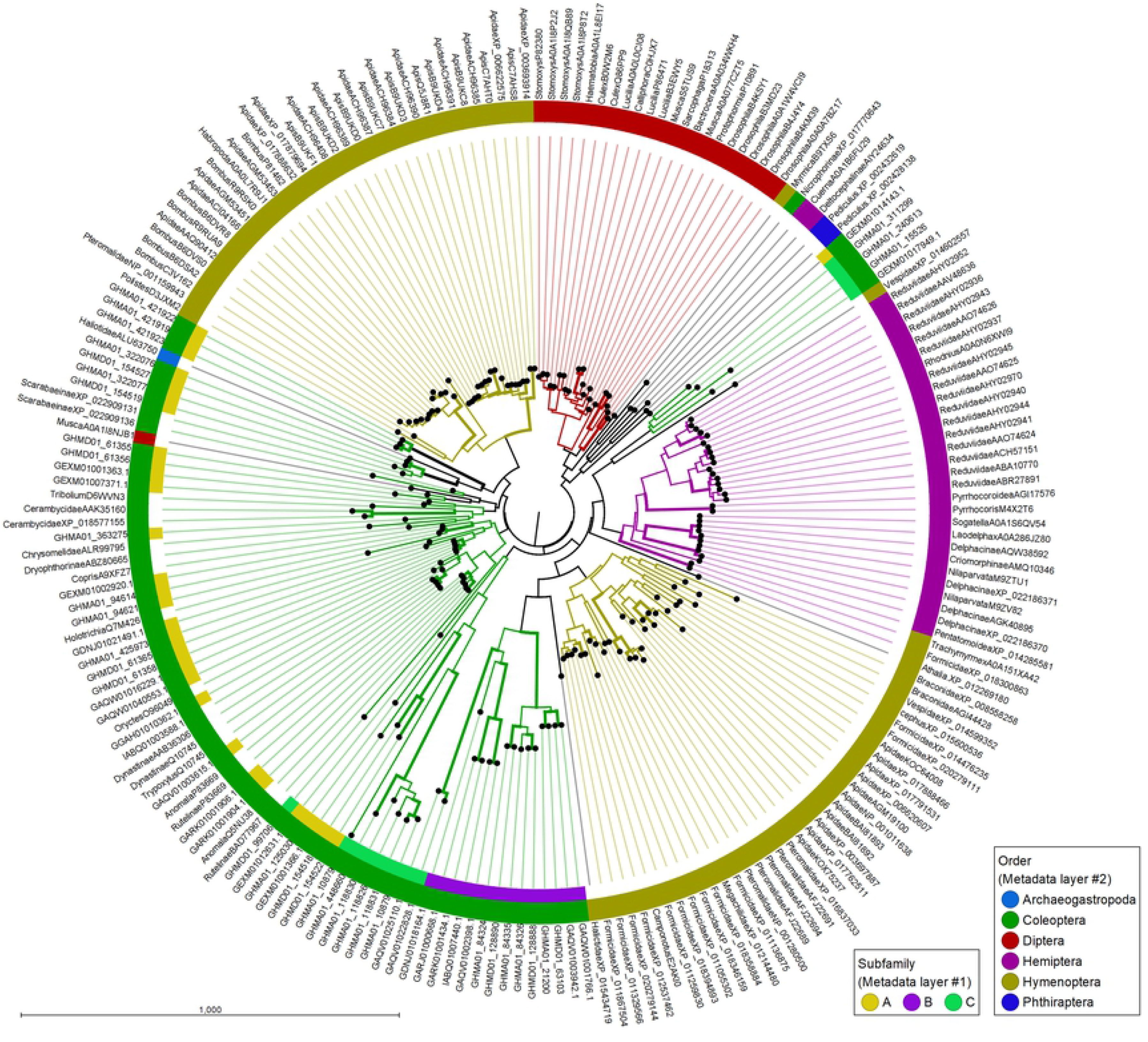
Neighbor-joining similarity dendrogram of invertebrate defensins. Sequences from the mature peptides were aligned by MUSCLE. The taxonomic distribution of the sequences and the corresponding defensin group is indicated in the color code presented. The confidence values of the branches were calculated with 10,000 bootstrap replicates. Thicker lines show branches with bootstrap threshold value > 70.

### Attacin family description

Attacins are peptides with molecular masses of 20-23 kDa with a wide spectrum of antimicrobial activities, including Gram-positive and Gram-negative bacteria, fungi, and protozoan parasites. As described for defensins, attacins are synthesized as pre-pro-proteins containing a signal peptide, and a conserved R-X-[RK]-R motif, which can be recognized by furin-like enzymes. The mature peptide is composed of an N-terminal attacin domain, followed by a glycine-rich segment. Attacins may be further divided according to basic and acid attacins [28,101–103].

In this study, we identified 41 new Scarabaeidae attacin sequences that fulfill the characteristics described (**Fig 9**). They are also recognized as sequences from this family by the Pfam signature Attacin-C (PF03769) [104,105]. The predicted mature sequences of the identified attacins can be further divided into two groups (named group A or B) according to the net charge (**Fig 10**). Compared to group B attacins, those belonging to group A are more cationic as they are highly enriched in positively charged residues. Additionally, group A attacins contain significantly more GNTS polar residues.

**Fig 9.**
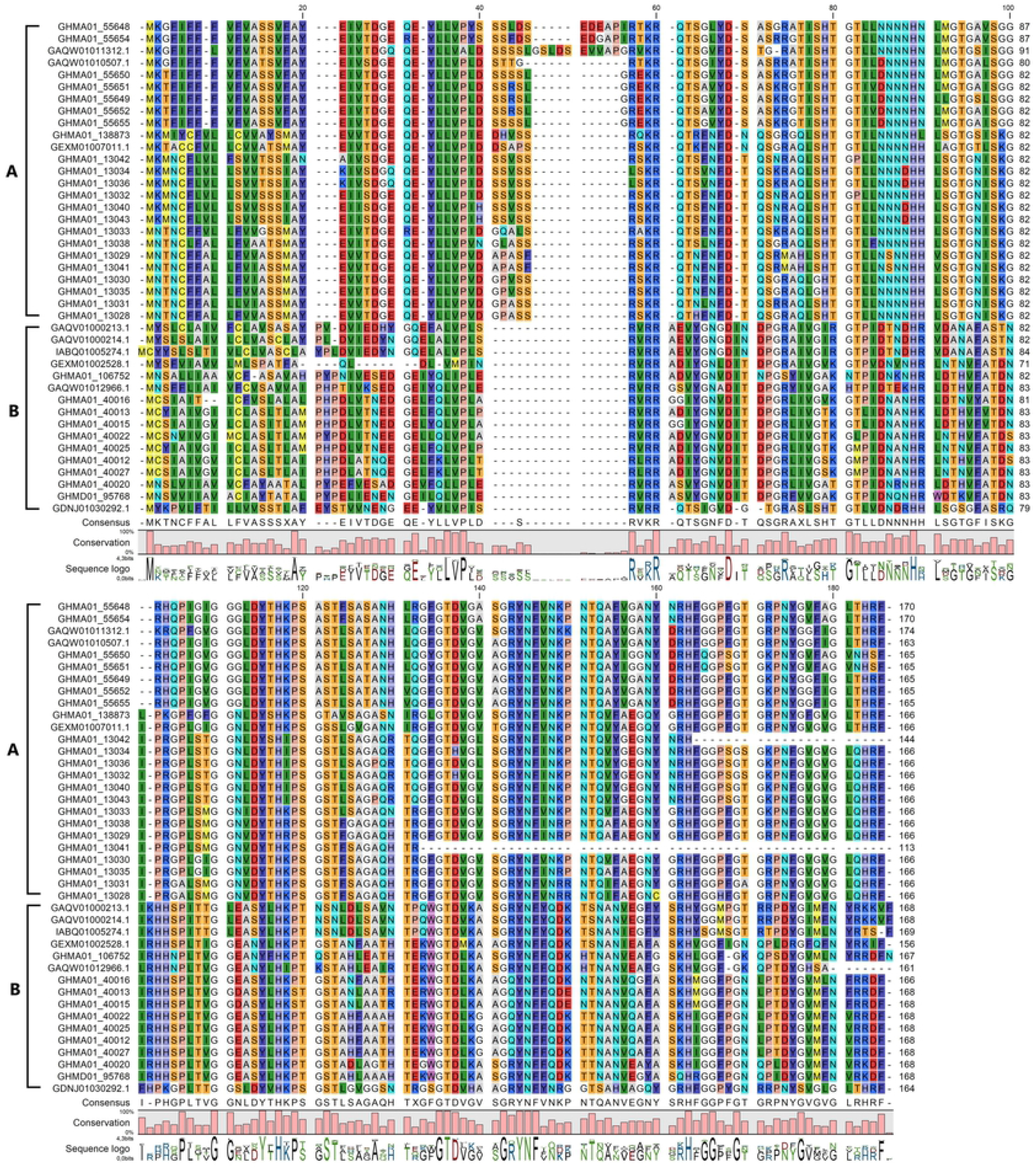
Multiple-sequence alignment of attacin HDP found in Scarabaeidae. Position 19 in the alignment represents the cleavage site of the signal peptide predicted with SignalP. The propeptide comprises positions 20 to 60. Position 80 indicates the propeptide furin-like cleavage site.

**Fig 10.**
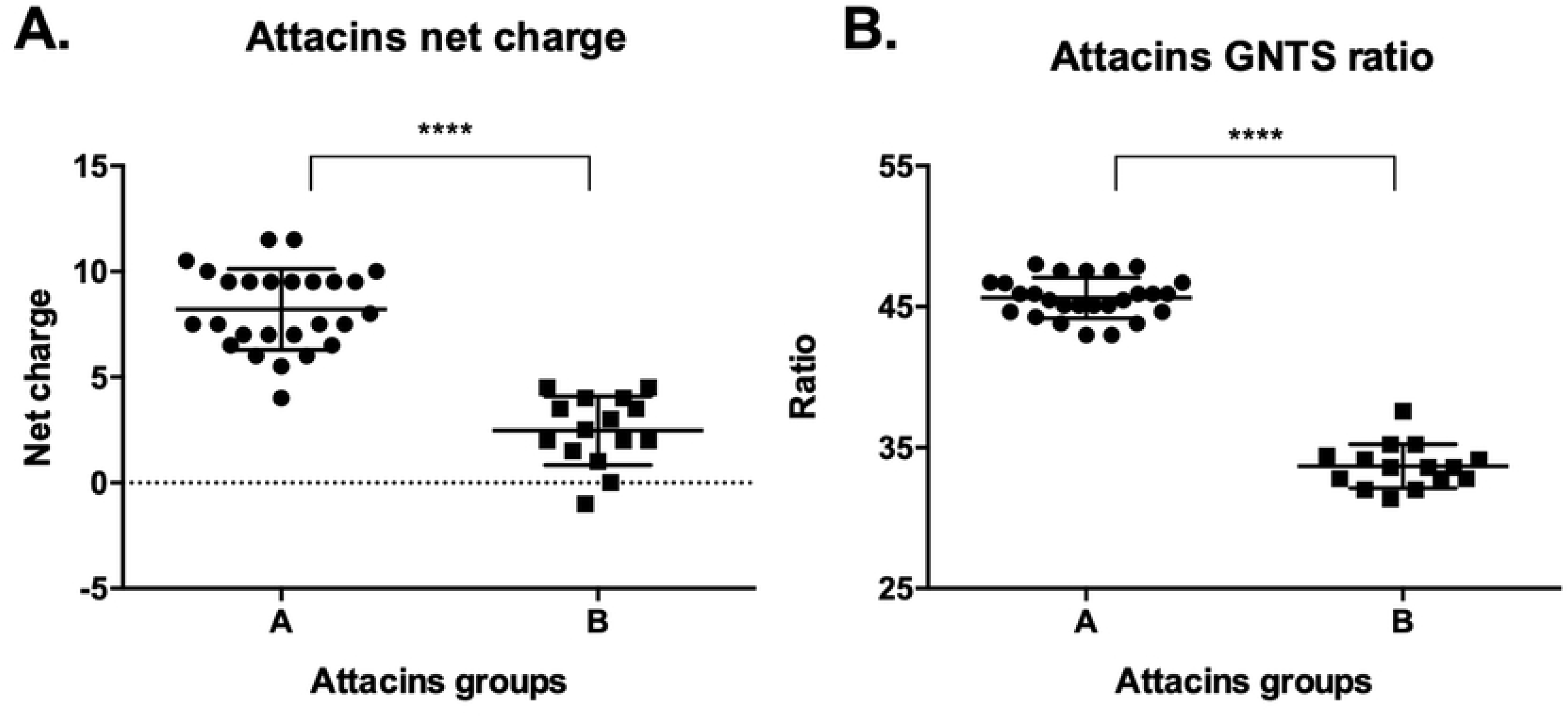
Physicochemical characteristics of the Scarabaeidae attacins. **A**. Net charge and **B**. GNTS composition. Statistical differences were evaluated by ANOVA test with Tukey’s multiple comparisons (p-value: ****<0.0001).

The secondary and tertiary structures predicted for Scarabaeidae attacins were characterized by a predominant antiparallel eight string beta-sheet configuration (**Fig 11**). The structural similarity was higher for group A attacins, as evidenced by higher TM scores (Group A= 0,73 and Group B= 0,58). The tertiary structure partially resembles those adopted by barrel channels. Interestingly, their structure was modeled by using the *E. coli* TamA barrel domain (PBD:4N74) as template by the automatic server predictor RaptorX. TamA forms a barrel channel with 16 transmembrane beta-sheets that translocates protein substrates across bacterial membranes [106].

**Fig 11.**
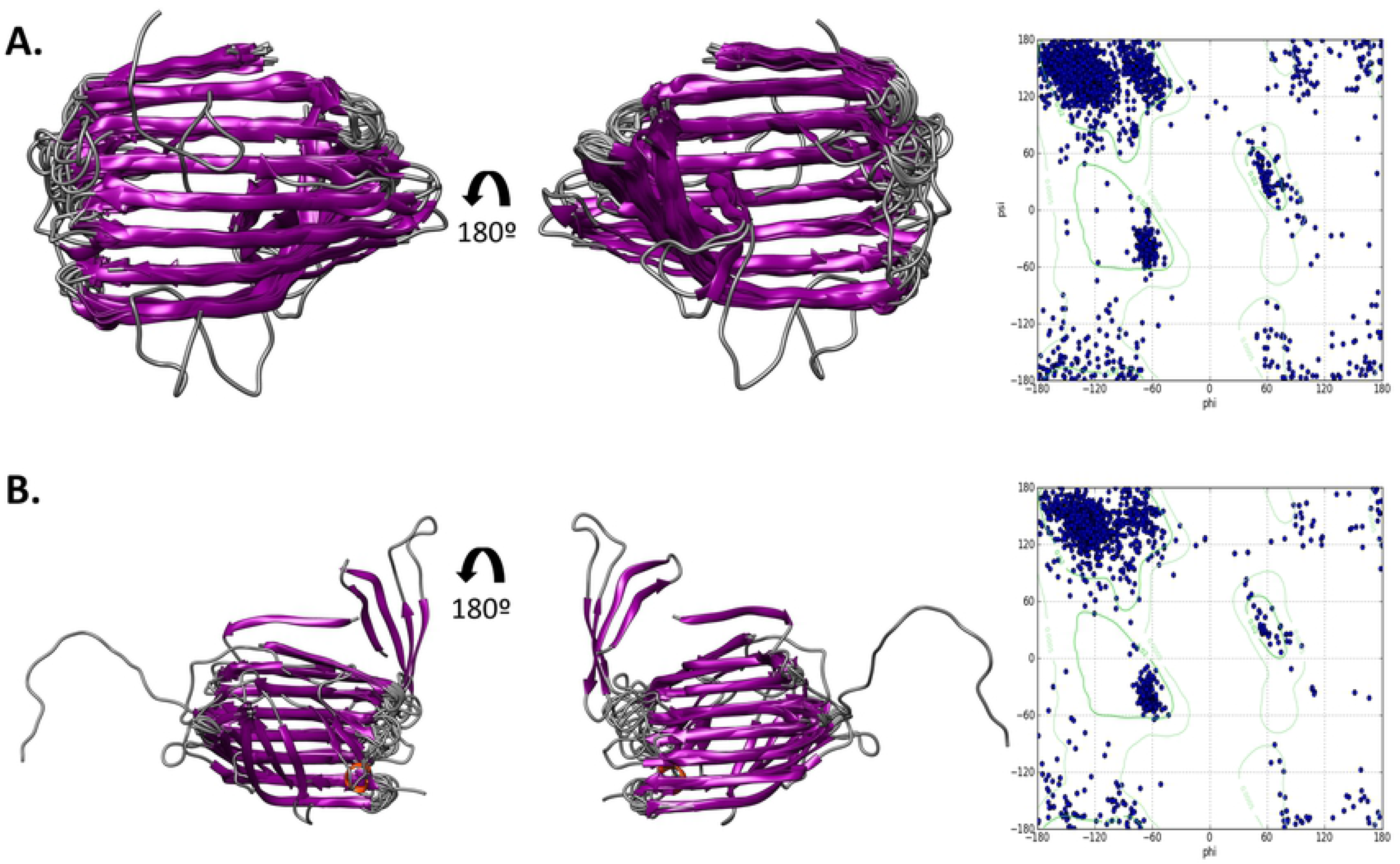
Structural superposition (left) and Ramachandran plot (right) of the mature attacins. **A**. Analysis of group A, TMscore=0.7366. **B**. Analysis of group B, TMscore=0.5844. The analysis was conducted in DeepAlign.

Based on these observations we hypothesized that attacin antimicrobial activity may be related to the formation of similar configurations in the bacterial membranes. To evaluate this possibility further, a protein-protein interaction modeling was conducted. This analysis predicts homodimers of attacins with barrel-like structures containing an outward hydrophobic face, conserved acidic residues facing inward and glycine-rich regions corresponding to the beta loops (**Fig 12**). These findings are compatible with the experimental results of attacin E of *Hyalophora cecropia*, which is targeted to the outer membrane of *E. coli* and facilitates the penetration of cations such as sodium and potassium [106]. The tertiary structure of attacins has not been experimentally determined in the PDB. The structural information available was obtained by circular dichroism (CD) on a recombinant attacin encoded by *Hyalophora cecropia*. An α-helical structure for this protein was deduced based on the presence of a single peak at 222 nm in CD [101]. However, the distinctive feature of α-helical proteins in CD are negative bands at 208 and 222 nm [107], so the structure of this protein family remains an open question.

**Fig 12.**
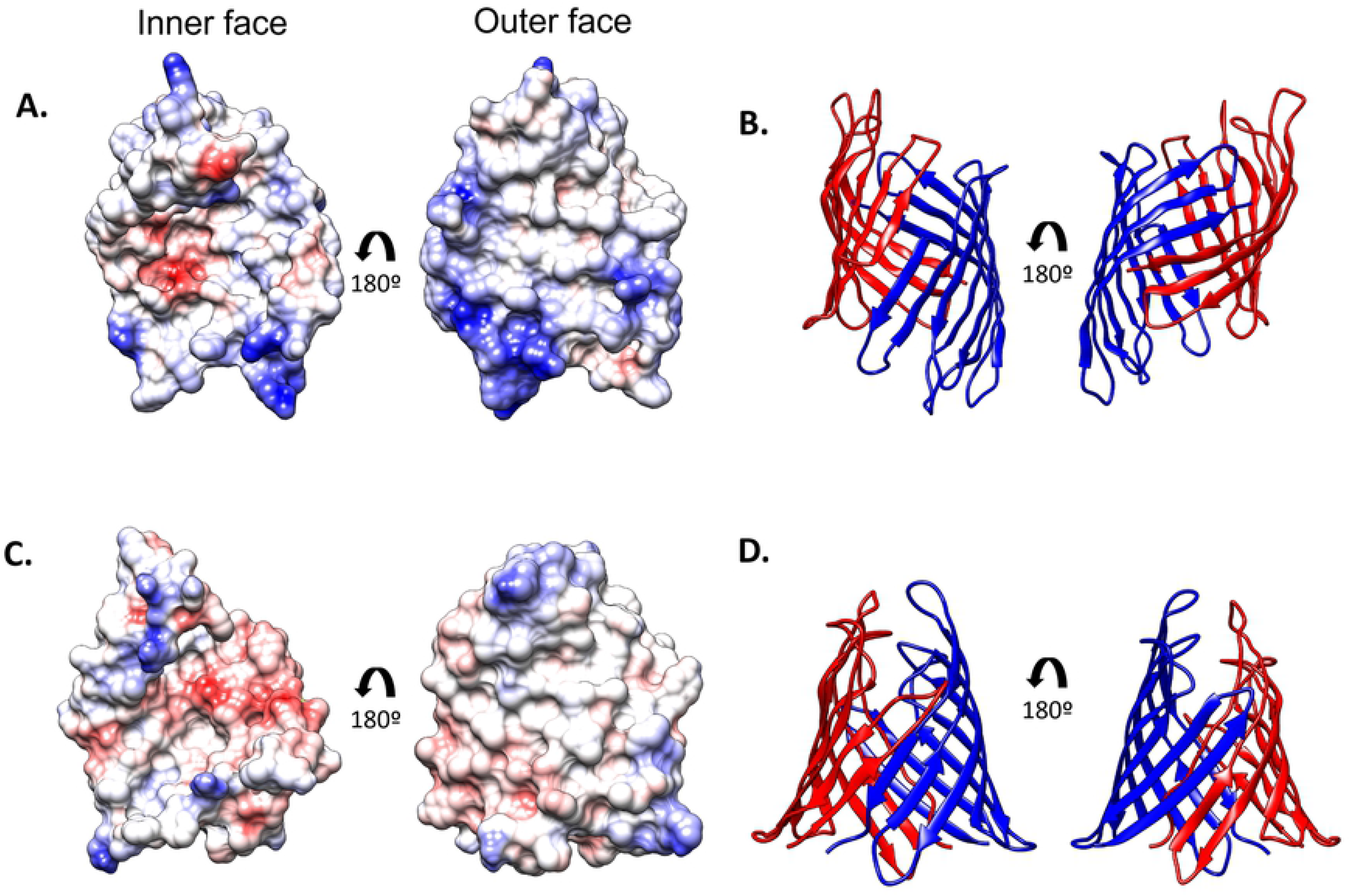
Attacin structural protein-protein interaction prediction. **A**. and **B**. Attacin A (GHMA01_13033). **C**. and **D**. Attacin B (GHMA01_106752). **A**. and **C**. Monomer electrostatic potential, according to Coulomb’s law surface coloring (red −10, white 0, blue 10 Kcal/(mole^e)). **B**. and **D**. Dimer protein-protein interaction prediction in ribbons chain coloring.

Dendrogram for insect attacins shows a clear overrepresentation of sequences from Diptera, especially flies representing 59% of the sequences. This group of diptera sequences representing flies forms a single clade, as well as Lepidoptera, for Coleoptera attacins; the group in a distant clade in close proximity to a second diptera group representing mosquitoes. The Coleoptera attacins had a clear differentiation in the A and B groups described. Only two sequences of Coleoptera attacins were previously annotated in the Interpro (Oryctes A0A0T6BE25; Oryctes A0A0T6BFF4) and were grouped into the B Scarabaeidae attacins (**Fig 13**).

**Fig 13.**
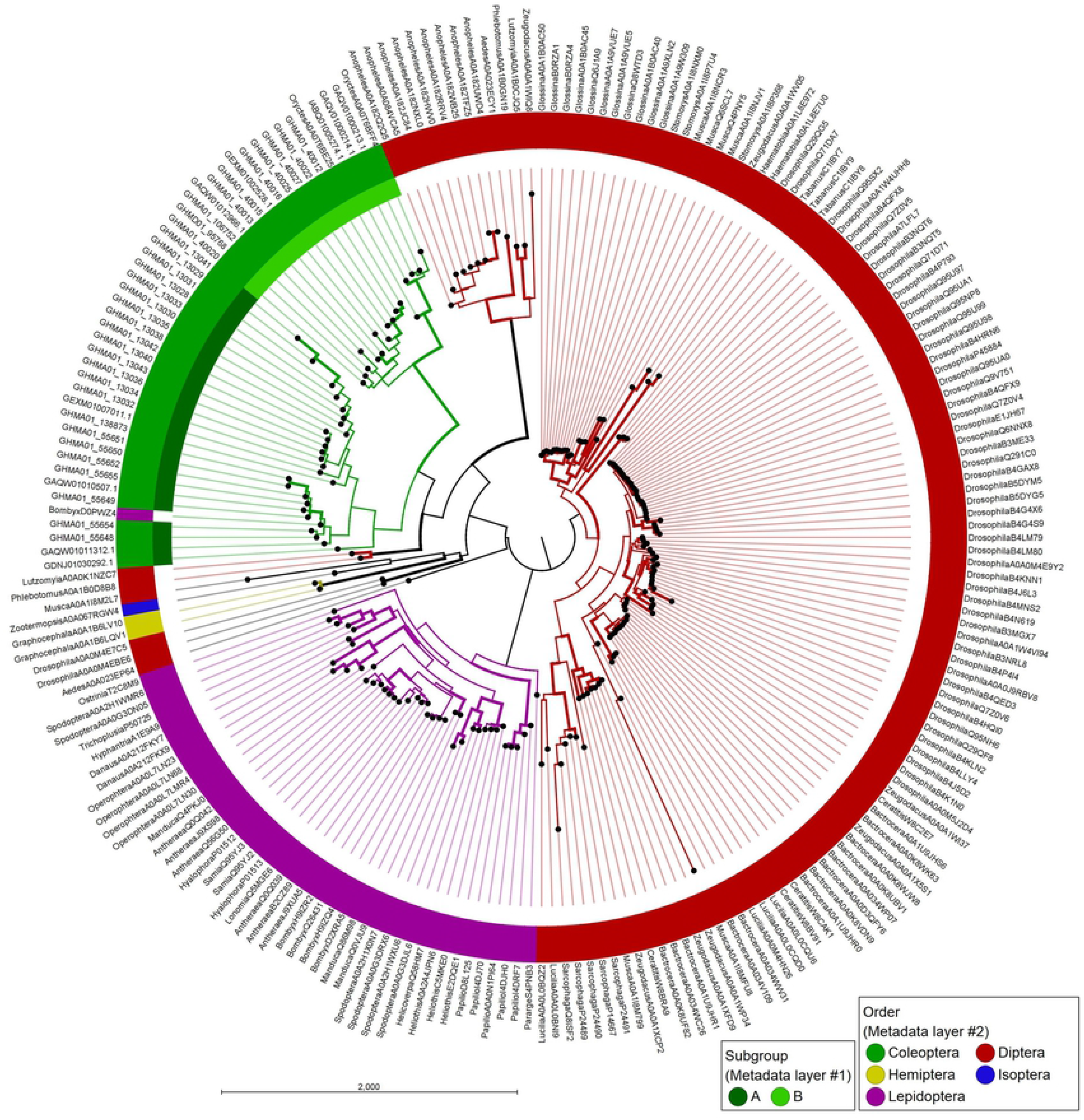
Neighbor-joining similarity dendrogram of insect attacins. Sequences from the mature peptides were aligned by MUSCLE. The taxonomic distribution of the sequences and the corresponding attacin group is indicated in the color code presented. The confidence values of the branches were calculated with 10.000 bootstrap replicates. Thicker lines show branches with bootstrap threshold value > 70.

### Coleoptericin family description

Coleoptericins are a group of antimicrobial peptides found exclusively in the Coleoptera order. The first coleoptericin was described in the tenebrionid beetle, *Zophobas atratus* [33]. This family is characterized by a signal peptide and a furin-like cleavage site with a mature peptide of 71 to 75 residues [108]. This group of HDP shows antimicrobial activity to Gram-positive and Gram-negative bacteria, with bacteriostatic and bactericidal activity at low and high concentrations respectively. Their action mechanism is unclear, but liposome leaking experiments suggest that it does not involve the formation of pores. Instead, they elicit the formation of an elongated and chain formation morphology in the bacteria [32]. A coleoptericin has recently been shown to play a central role in host-symbiont interactions in the weevil *S. zeamais* [50].

Coleoptericins encoded by Scarabaeidae (37 sequences) were identified as pre-pro-proteins with a furin-like cleavage site, they share the motif G-P-[GNS]-[KR]-[GSA]-K-P from position 97 to 103 (**Fig 14**). They were classified according to their amino acid sequence conservation into three major groups: 1. Group A (18 sequences) with approximately 73 amino acid residues in length, a cationic character with a mean net charge of 5.63, and a relative disorder structure with a random coil-beta sheet structure (TM score: 0.27) (**Figs 15–16**); 2. Group B is a small group in Scarabaeidae (three sequences), but shows high similarity with already reported coleoptericins (**Fig 17**). They are typically 72 residues in length, with a mean net charge of 5 and a coil-beta sheet-helix-coil configuration in the tertiary structure (**Fig 16**); 3. Group C (17 sequences) is 57 to 59 amino acids in length and is enriched in acidic residues that provide an mean positive net charge of 1.41. They have a relatively disordered structure with a coil-beta sheet-helix-coil configuration (**Figs 14–17**). In general, the conserved coleoptericin motif is structurally preceded by an alpha helix or related to the third beta loop in the structural conformation.

**Fig 14.**
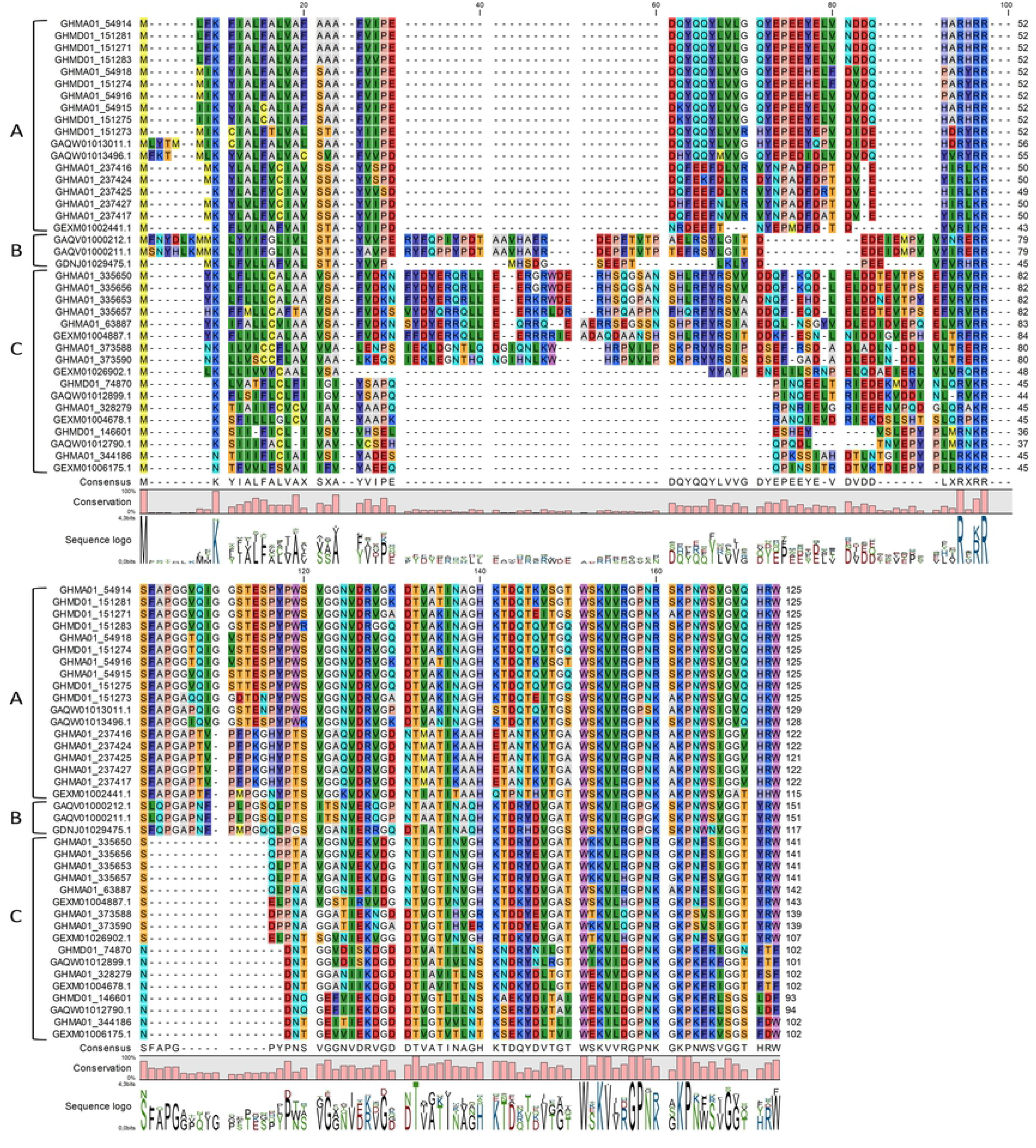
Multiple sequence alignment of Coleoptericin HDP found in Scarabaeidae. Position 23 in the alignment represents the cleavage site of the signal peptide predicted with SignalP. Positions 94 to 97 furin-like cleavage site.

**Fig 15.**
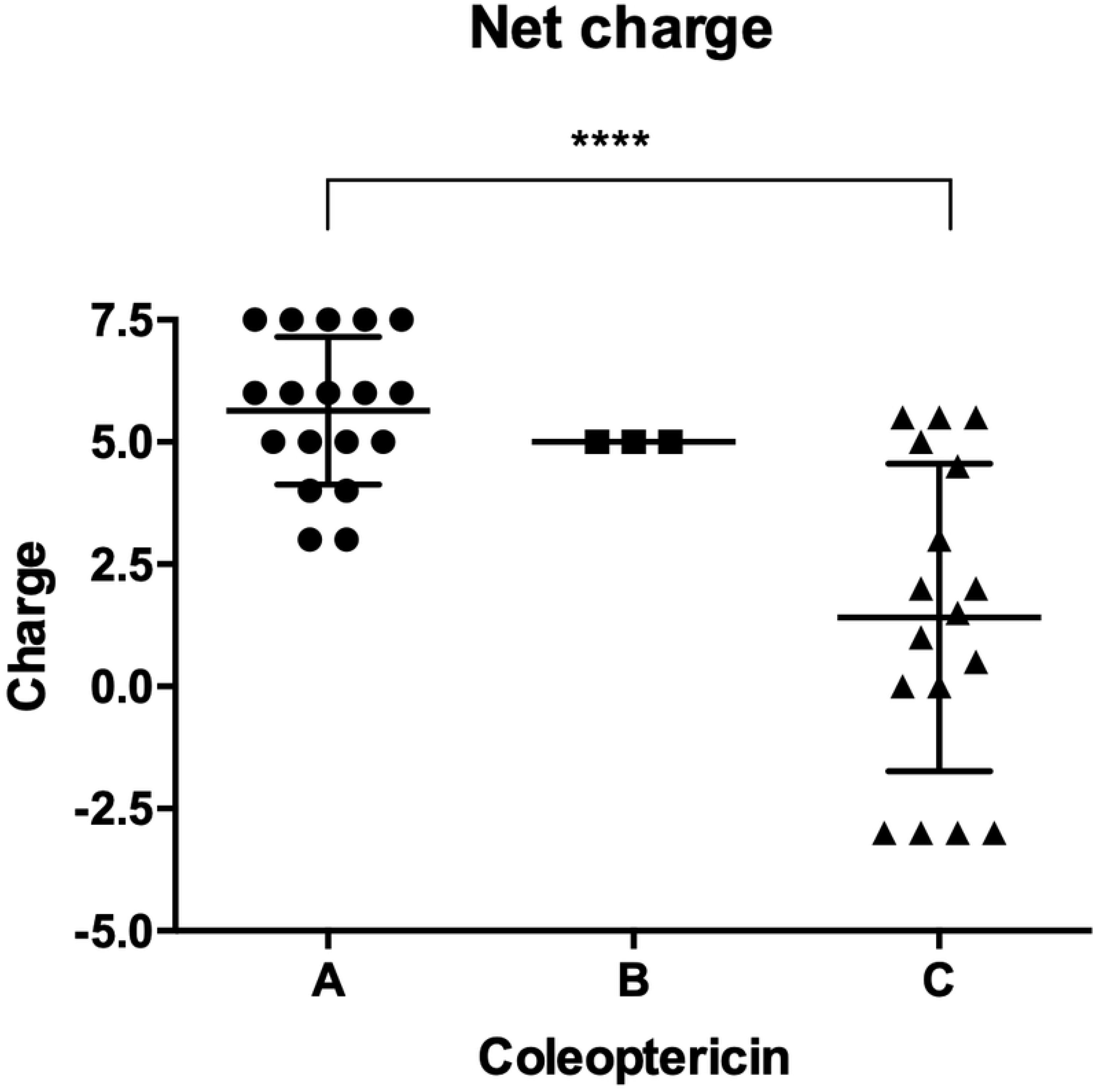
The net charge of the Scarabaeidae Coleoptericins. Statistical differences were evaluated by ANOVA test with Tukey’s multiple comparisons (p-value: ****<0.0001).

**Fig 16.**
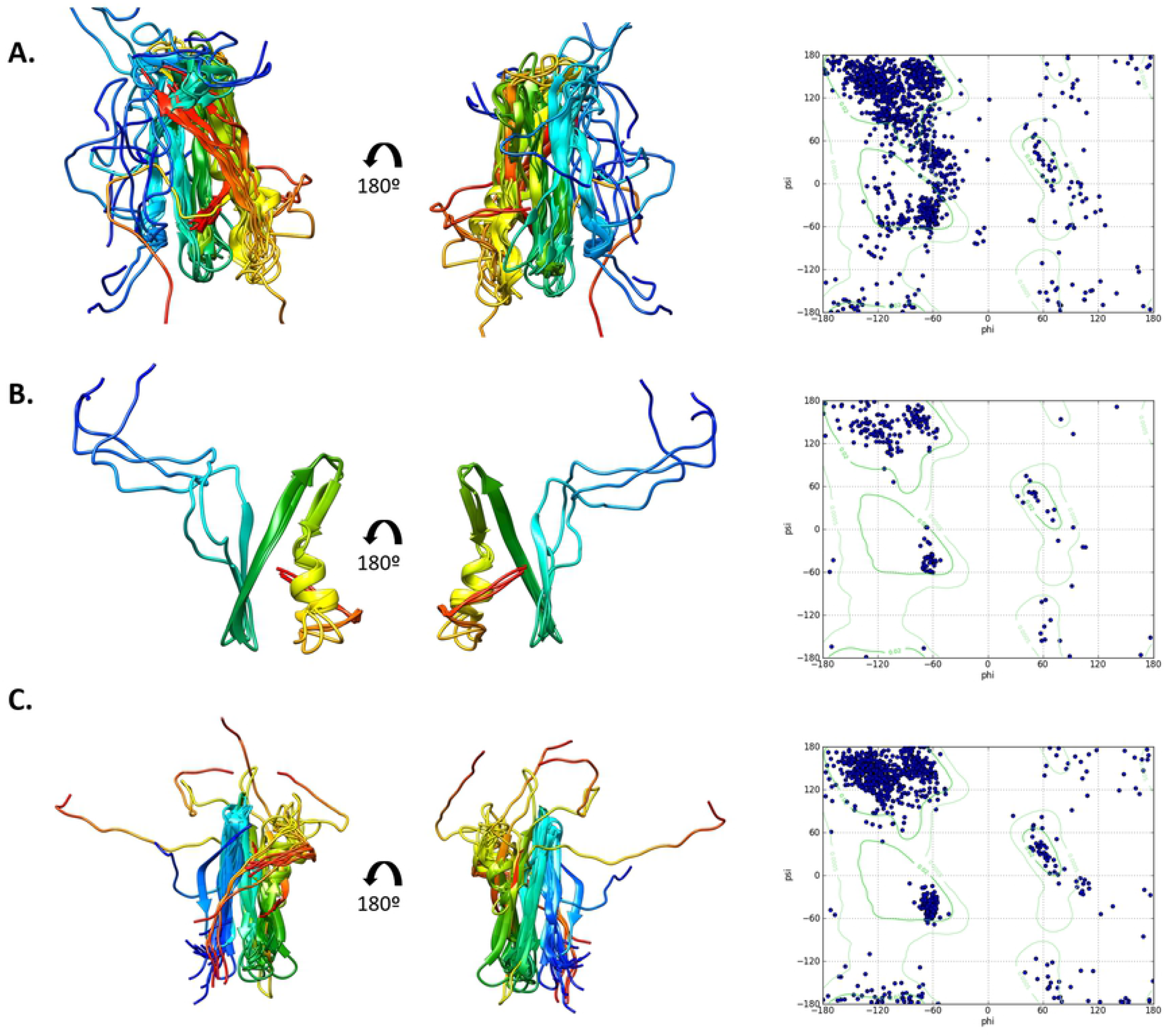
Structural superposition (left) and Ramachandran plot (right) of the mature coleoptericins. **A**. Analysis of group A, TMscore=0.2753. **B**. Analysis of group B, TMscore=0.6248, **C**. Analysis of group C, TMscore=0.3925. The analysis was conducted in DeepAlign.

**Fig 17.**
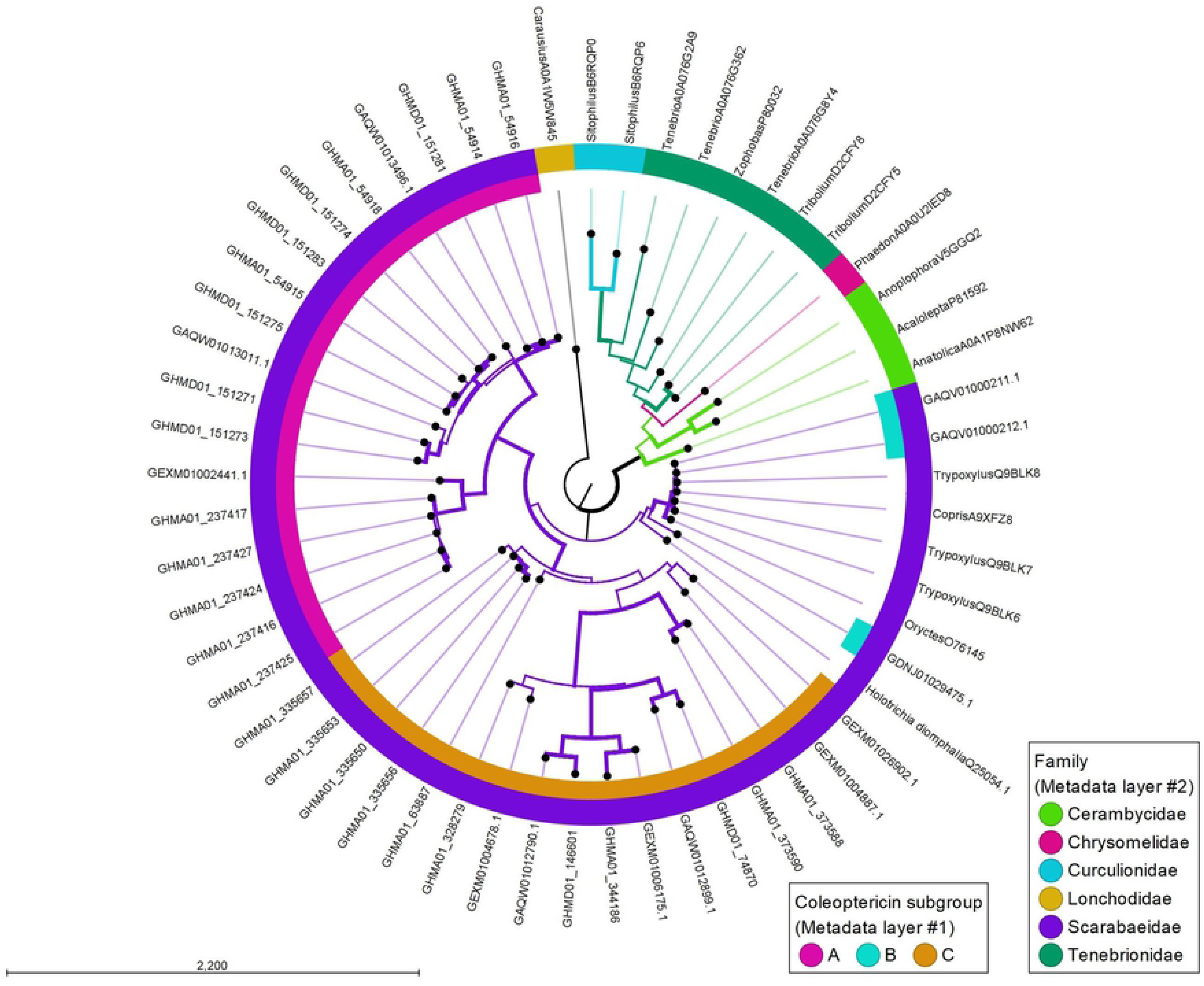
Neighbor-joining similarity dendrogram of Coleoptericins. The taxonomic distribution of the sequences and the corresponding coleoptericin group is indicated in the color code presented. A sequence from *Carausius* (Phasmatodea order) was used as the root node. The remaining sequences belong to the Coleoptera order. The confidence values of the branches were calculated with 10,000 bootstrap replicates. Thicker lines show branches with bootstrap threshold value > 70.

Coleoptericins A, B, and C identified form three separate groups exclusive of the Scarabaeidae family. This work, reports two new groups, A and C coleoptericins, which are a group in a different clade and are exclusive of Scarabaeidae (**Fig 17**).

### Distinct physicochemical properties characterize the HDP families of Scarabaeidae

Families of insect antimicrobial peptides have distinct physicochemical properties, amino acid composition, and structural features. This has allowed the classification of HDP according to them [13]. Insect antimicrobial peptides can be divided into four classes: α-helical peptides (cecropin and moricin), cysteine-rich peptides (insect defensin and drosomycin), proline-rich peptides (apidaecin, drosocin, and lebocin) and glycine-rich peptides (attacin and gloverin) [37,109]. With this in mind, those features were analyzed in the HDP identified from Scarabaeidae. Attacins and coleoptericins were the families with higher molecular mass (Mean: 12931Da SD: 1075Da and Mean: 7097Da SD: 736.6Da, respectively) and lowest hydrophobic nature (0.27 and 0.28 ratios respectively) (**Fig 18**). Defensins and cecropins showed similar hydrophobic ratios (0.37 and 0.47, respectively). As described for other members of cecropins [26], those identified in Scarabaeidae are characterized by high net charges (Mean: 9.25; SD: 1.71) and pI values (Mean: 11.3; SD: 0.41) (**Fig 18**). These characteristics are reflected in the amino acid composition of the peptide families (**Fig 19**). Cecropins show a higher proportion of positively charged amino acids, and attacins and coleoptericins a higher content of polar residues. Prolines and glycines are structurally important amino acids, hence, differences were sought in the content of these amino acids in the HDP. In contrast to the other two families, more than 10 % of the amino acids encoded by attacins and coleoptericins correspond to these two residues (**Fig 19**).

**Fig 18.**
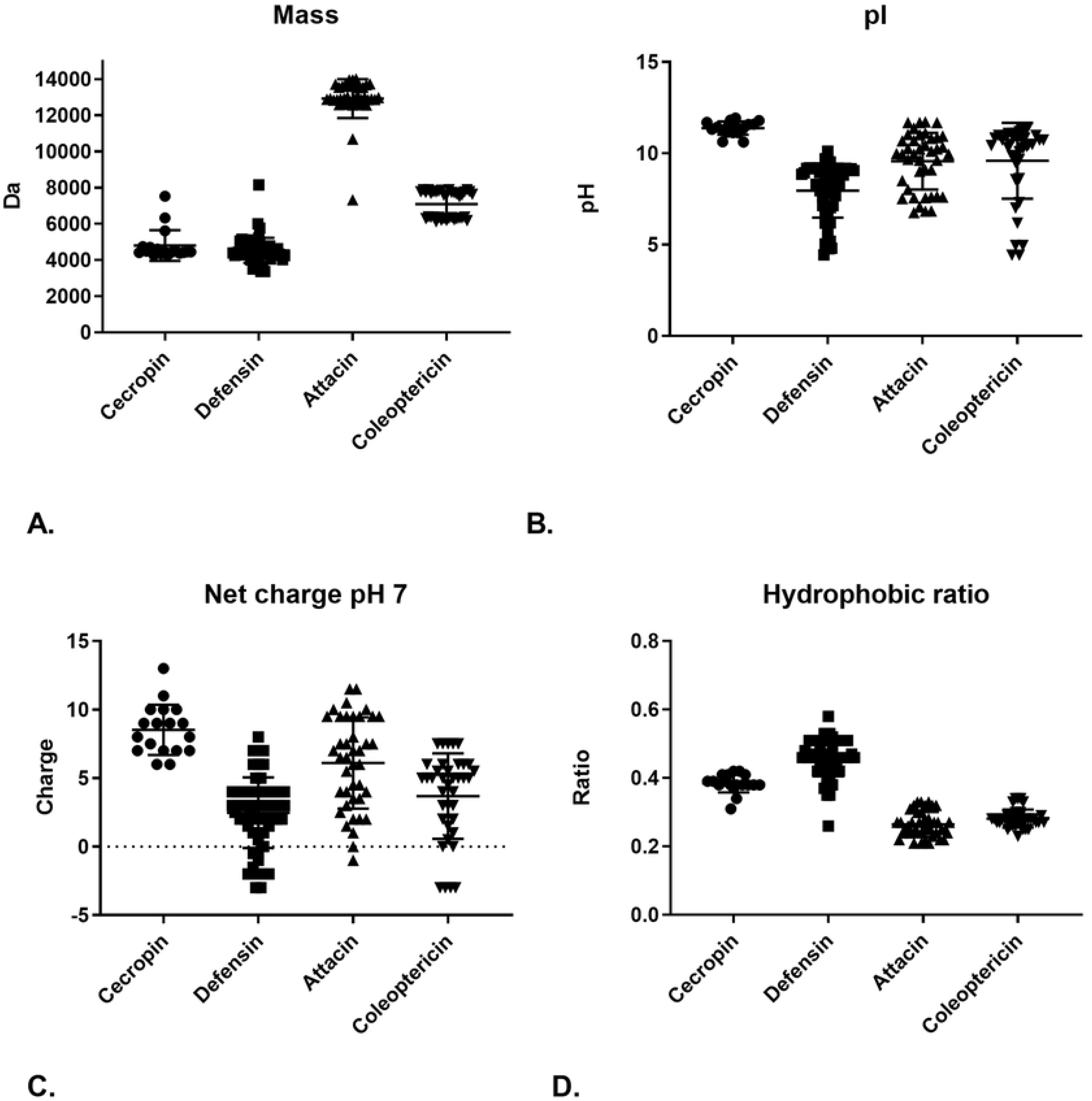
Physicochemical properties of Scarabaeidae HDP families predicted with APD3. **A**. Molar mass. **B**. Isoelectric point. **C**. Net charge. **D**. Hydrophobic ratio

**Fig 19.**
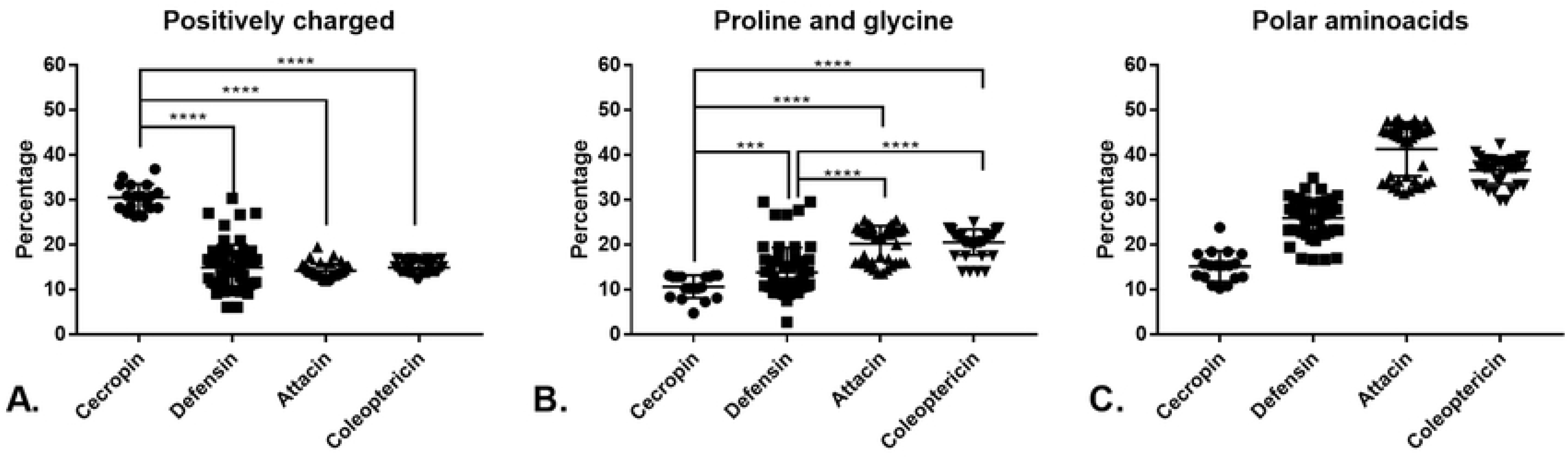
Representation of amino acid groups in the families of Scarabaeidae HDP. **A**. Positively charged amino acids (RK). **B**. Proline and glycine amino acids **C**. Polar amino acids (GNST), statistical difference was observed among all groups (****). Statistical differences were evaluated by ANOVA test with Tukey’s multiple comparisons (p-value: ***<0.001; ****<0.0001).

The amino-acid distribution represents a key characteristic to classify HDP families, like proline, lysine, and cysteine-rich antimicrobial peptides [13,37,109,110]. According to this, in Scarabaeidae HDP it was found that positively charged amino-acid proportion only differentiates the cecropin family (**Fig 19A**). The percentage of glycine and proline obtained showed a statistical difference among all groups except between the attacin and coleoptericin. The polar amino acid (GNST) distribution differentiates the HDP families with a statistically significant difference (**Fig 19B, C**).

### Antimicrobial activity prediction of dung beetle HDP *in silico*

Development of sequence-based computational tools can be helpful in identifying candidate HDP for experimental characterization. Different tools classified the sequences according to different parameters, like physicochemical characteristics, compositional amino acids, family signatures, and hidden Markov models (HMM); with complementary approaches regarding the accuracy, capacity of prediction, and training set for the construction of the program [13,68,69].

The cecropin family in CAMPr3 had 17 of 18 sequences with a positive prediction with four different tools (Random forest, support vector machine, discriminant classifier, artificial neural network) compared to iAmPred which shows lower scores. iAmPred and Campr3 predicted high antimicrobial probability to defensins (groups A and B) and attacins; and low antimicrobial probability to cecropins and coleoptericins. ClassAmp2 predicted a high probability for all the families involved (**Fig 20**). This high probability prediction for defensins and low activity prediction for cecropins may be related with the fact that β-defensins had been reported in a wide range of organisms and thus have a higher diversity and representation in the data bases and training libraries, whereas, cecropins had been found majorly in insects (limited taxon) with a low level of representation for some groups, like Coleoptera [111].

**Fig 20.**
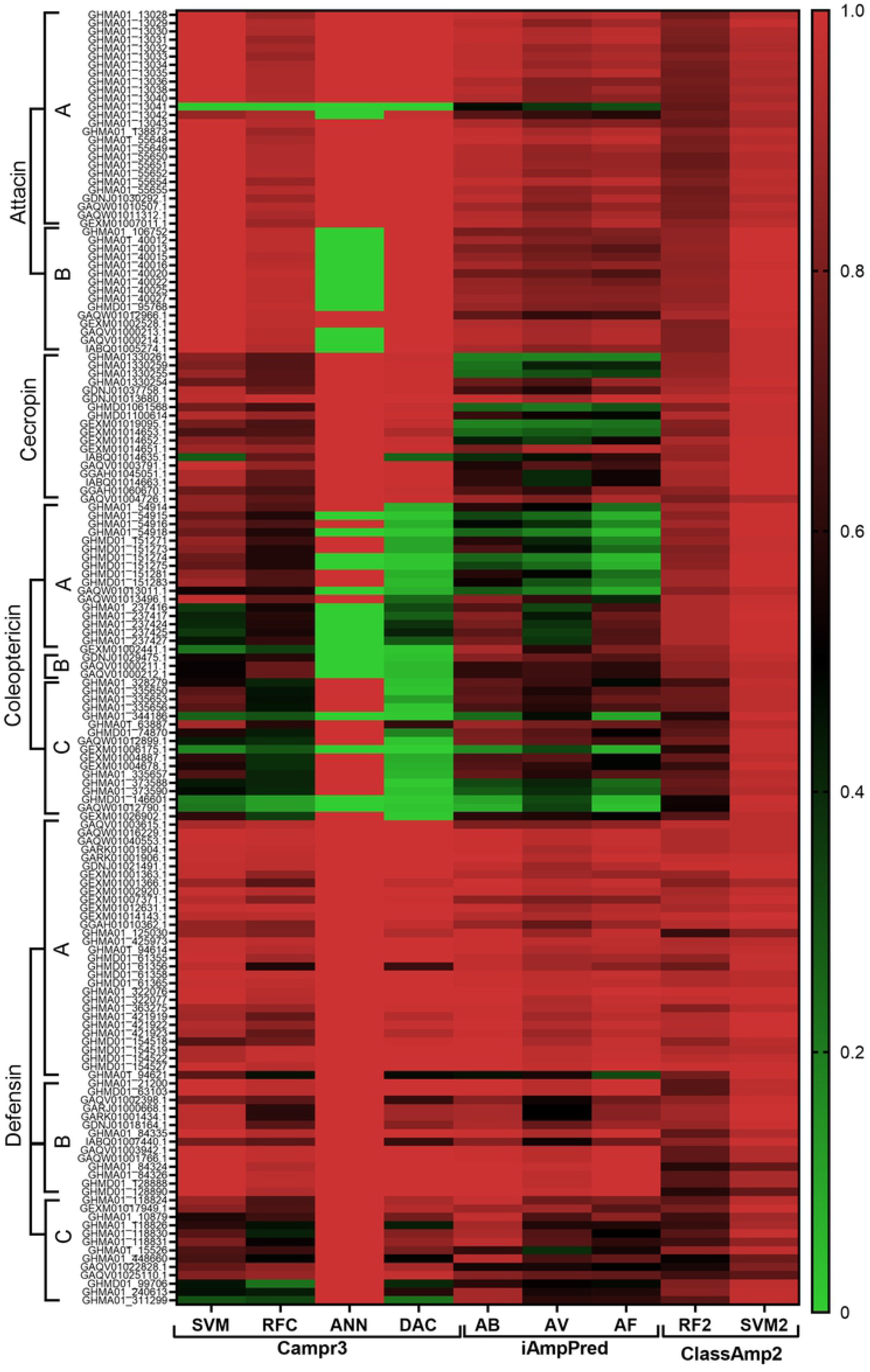
Heat map for the antimicrobial prediction of Scarabaeidae HDP. The gradient represents the probability score for each server from inactive (green), medium activity (black), and active (red). The HDP families were separated in their respective groups: Attacin A and B; Coleoptericins A, B, and C; Defensins A, B, and C. SVM: support vector machine; RFC: Random forest; ANN: artificial neural network; DAC discriminant analysis classification; AB: antibacterial; AV: antiviral; AF: antifungal.

In general, CAMPr3 has a higher probability of prediction compared with iAMPpred. These results may be due to the greater data set used to construct the prediction in CAMPr3, including family signatures, compared with iAMPred; nevertheless, iAMPpred includes a wide diversity of physicochemical parameters and compositional AA parameters. The cecropin family has a lower probability in iAMPpred, compared with CAMPr3; this result may be explained by the underrepresentation of the cecropin family of Coleoptera in the databases and the relative divergence of the Scarabaeidae cecropins compared with other taxa (**Fig 4**). This also may explain the low prediction probabilities for the coleoptericins in general.

## Conclusion

Using HDP has been suggested as a complement to traditional antibiotics, and insects are considered attractive sources of novel HDP, given that their immune response relies – in part – on the expression of diverse families of these peptides showing strong antimicrobial activity at low concentrations and against a wide range of pathogens. So far, identification of insect HDP has focused on the Hemiptera, Hymenoptera, Lepidoptera, and Diptera orders [85]. In Coleoptera, the most diverse order of insects, only a few HDP have been reported [2,24,53,85]. This work identified 155 novel sequences of HDP found in nine transcriptomes from seven Coleoptera species: *D. satanas* (n= 76; 49.03%), *O. curvicornis* (n= 23; 14.83%), *T. dichotomus* (n= 18; 11.61%), *O. nigriventris* (n= 10; 6.45%), *Heterochelus* sp. (n= 6; 3.87%), *O. conspicillatum* (n= 18; 11.61%) and *P. japonica* (n= 4; 2.58%). Two information sources were retrieved: 1) *De novo* transcriptomic data from two neotropical Scarabaeidae species (*Dichotomius satanas* and *Ontophagus curvicornis*); 2) Sequence data deposited in public sequence databases. The HDP sequences from Coleoptera were identified based on similarity to known HDP insect families. New members of defensins (n = 58; 37.42%), cecropins (n = 18; 11.61%), attancins (n = 41; 26.45%), and coleoptericins (n = 38; 24.52%) were detected. These families were described based on their physicochemical properties, structural features, and sequence similarity. Both novel and previously described attributes were observed in different families. These features are summarized in **Fig 21**. Most HDP sequences show predicted antibacterial, antiviral, and antifungal activities (**Fig 20**), so they may be promising candidates for experimental characterization in anti-microbial and cytotoxicity screening assays. Additionally, this work would also help distinguish key attributes associated with different peptide activities, allowing identification and description of new anti-microbial peptides.

**Fig 21.**
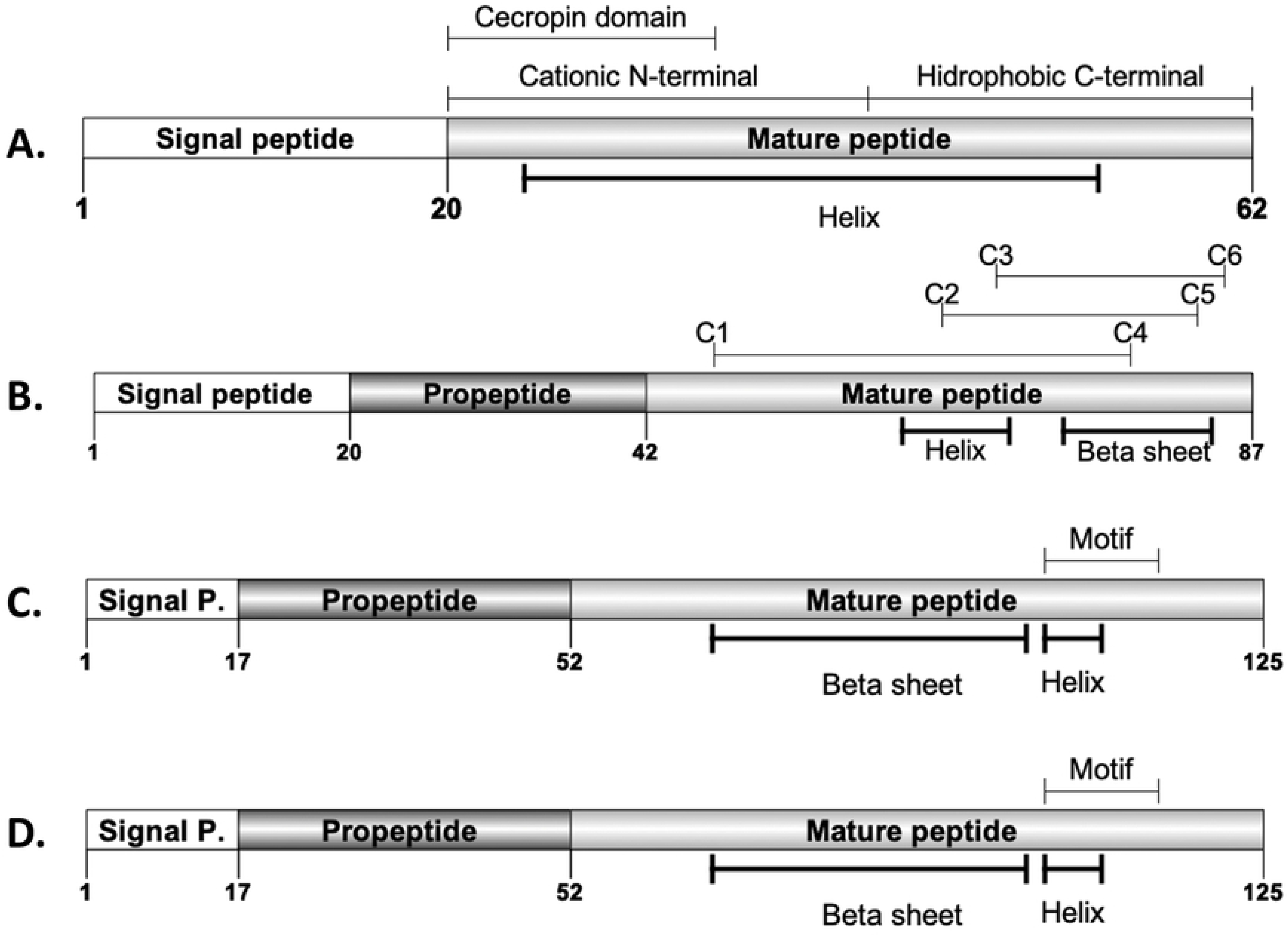
Scarabaeidae HDP general sequence and structural features **A**. Cecropins. **B**. Defensins. **C**. Attacins **D**. Coleoptericins.

## Acknowledgments

Students of the research seedbed 2017, Universidad del Quindío, Jean Carlos Sanchez Rojas, Luz Dahiana Erazo Urquijo; Alejandra Vélez García; Helen Vanessa Pinchao Salgar and Angélica Maria Madrid Ortiz.

